# Structural Basis of ECF-σ-Factor-Dependent Transcription Initiation

**DOI:** 10.1101/381020

**Authors:** Wei Lin, Sukhendu Mandal, David Degen, Min Sung Cho, Yu Feng, Kalyan Das, Richard H. Ebright

## Abstract

Extracytoplasmic (ECF) σ factors, the largest class of alternative σ factors, are related to primary σ factors, but have simpler structures, comprising only two of the six conserved functional modules present in primary σ factors: region 2 (σR2) and region 4 (σR4). Here, we report crystal structures of transcription initiation complexes containing *Mycobacterium tuberculosis* RNA polymerase (RNAP), *M. tuberculosis* ECF σ factor σ^L^, and promoter DNA. The structures show that σR2 and σR4 of the ECF σ factor occupy the same sites on RNAP as in primary σ factors, show that the connector between σR2 and σR4 of the ECF σ factor--although unrelated in sequence--follows the same path through RNAP as in primary σ factors, and show that the ECF σ factor uses the same strategy to bind and unwind promoter DNA as primary σ factors. The results define protein-protein and protein-DNA interactions involved in ECF-σ-factor-dependent transcription initiation.

## INTRODUCTION

Bacterial transcription initiation is carried out by an RNA polymerase (RNAP) holoenzyme comprising RNAP core enzyme and σ factor (reviewed in Feklistov et al., 2014). Bacteria contain a primary σ factor (group-1 σ factor; σ^70^ in *Escherichia coli;* σ^A^ in other bacteria) that mediates transcription initiation at most genes required for growth under most conditions and sets of alternative σ factors that mediate transcription initiation at sets of genes required in certain cell types, developmental states, or environmental conditions.

Group-1 σ factors contain six conserved functional modules: σ regions 1.1, 1.2, 2, 3, 3/4 linker, and 4 (σR1.1, σR1.2, σR2, σR3, σR3/4 linker, and σR4; Fig. 1A; reviewed in Feklistov et al., 2014). σR1.1 plays a regulatory role, inhibiting interactions between free, non-RNAP-bound, σ and DNA. σR1.2, σR2, σR3, and σR4 play roles in promoter recognition. σR2 and σR4 recognize the promoter −10 element and the promoter −35 element, respectively, and σR1.2 and σR3 recognize sequences immediately downstream and immediately upstream, respectively of the promoter −10 element. The σR3/4 linker plays multiple crucial roles (Murakami et al., 2002; Vassylyev et al, 2002; Mekler et al., 2002; Kulbachinskiy and Mustaev, 2006; Zhang et al., 2012; Basu et al., 2014; Pupov et al., 2014; Duchi et al., 2016; Lerner et al., 2016; Dulin et al., 2018). The σR3/4 linker connects σR2 to σR4; the σR3/4 linker enters the RNAP active-center cleft, where it interacts with template-strand ssDNA of the unwound “transcription bubble,” pre-organizing template-strand ssDNA to adopt a helical conformation and to engage the RNAP active center, thereby facilitating initiating-nucleotide binding and *de novo* transcription initiation; and the σR3/4 linker exits the RNAP active-center cleft by threading through the RNAP RNA exit channel. Before RNA synthesis takes place, the σR3/4 linker serves as a “molecular mimic” of RNA, or “molecular placeholder” for RNA, through its interactions with template-strand ssDNA and the RNAP RNA exit channel. As RNA synthesis takes place, the σR3/4 linker then is displaced, off of template-strand ssDNA and out of the RNAP RNA exit channel, driven by steric interactions with the 5’-end of the nascent RNA. The σR3/4 linker must be displaced from template-strand ssDNA during initial transcription; this requirement imposes energy barriers associated with initial-transcription pausing and abortive initiation. The σR3/4 linker must be displaced from the RNAP RNA exit channel during the transition between initial transcription and transcription elongation; this requirement imposes energy barriers that are exploited to trigger promoter escape and to transform the transcription initiation complexes into the transcription elongation complex.

**Figure 1.**
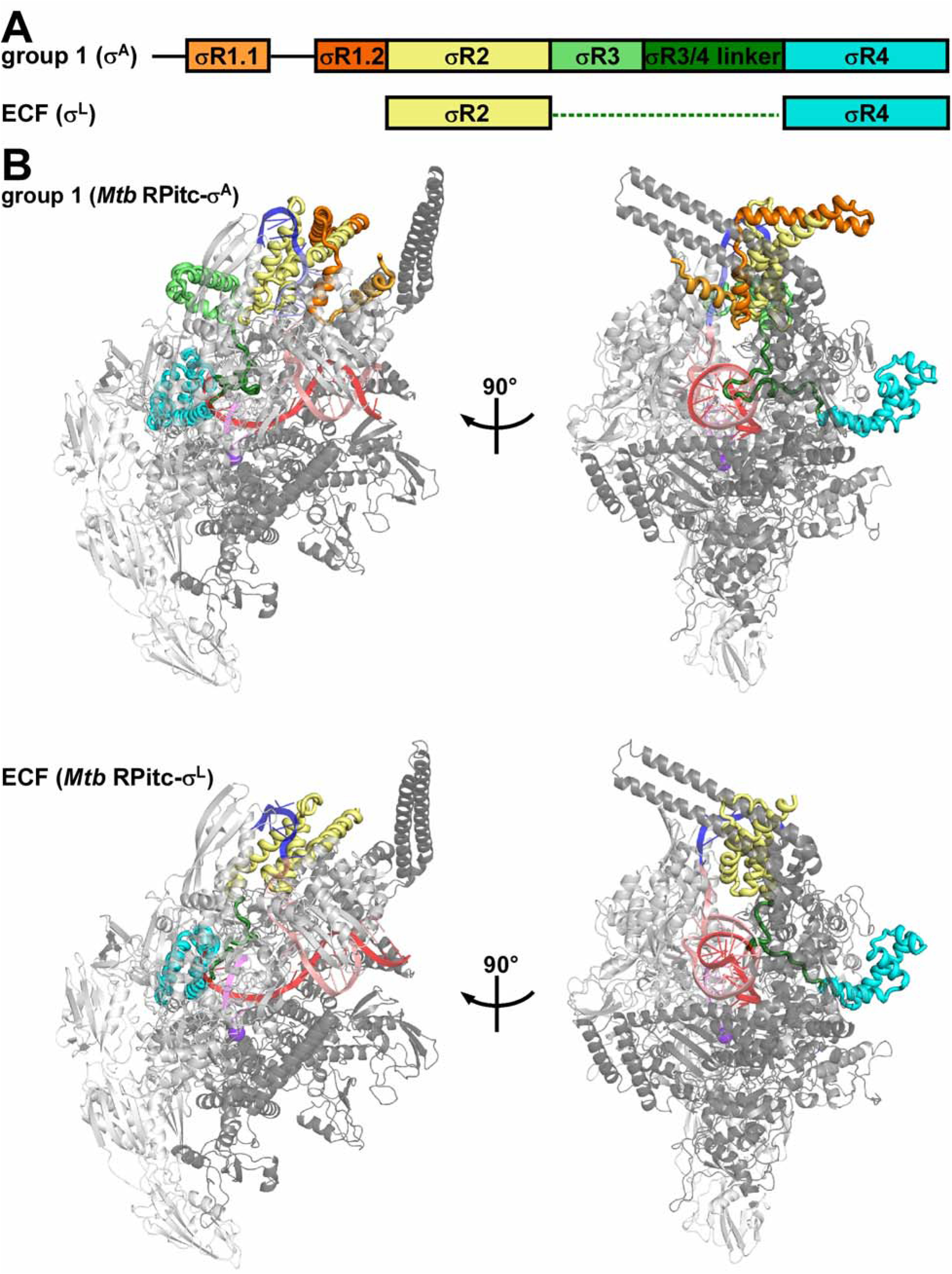
Structures of group-1 and ECF σ factors. **(A)** Structural organization of group-1 (σ^A^) and ECF (σ^L^) σ factors. Conserved regions σR1.1, σR1.2, σR2, σR3, σR3/4 linker, and σR4 are in pink, red, yellow, green, dark green, and cyan, respectively. Dashed line, non-conserved σR2/4 linker present in ECF σ factors. **(B)** Crystal structures of group-1 *(Mtb* RPitc-σ^A^) and ECF *(Mtb* RPitc-σ^L^) transcription initiation complexes (two orthogonal views of each). σ factors are shown in tube representations with conserved regions colored as in **(A)**. Gray ribbon, RNAP core enzyme; blue, pink, red, and magenta ribbons, −10 element of DNA nontemplate strand, rest of DNA nontemplate strand, DNA template strand, and RNA product; violet sphere, RNAP active-center Mg^2^+. Other colors are as in **(A)**. See Figs. S1, S2, and S7.

Crystal structures of RNAP holoenzyme and transcription initiation complexes containing group-1 σ factors define the protein-protein and protein-nucleic acid interactions involved in group-1-σ-factor-dependent transcription initiation; and extensive biochemical and biophysical characterizations define the protein-protein and protein-nucleic acid interactions and mechanisms involved in group-1-σ-factor-dependent transcription initiation (Murakami et al., 2002, 2013; Vassylyev et al, 2002; Zhang et al., 2012, 2014; Bae et al., 2013, 2015; Basu et al., 2014; Zuo and Steitz, 2015; Feng et al., 2016; Hubin et al., 2017; Lin et al., 2017, 2018).

Alternative σ factors--with the exception of the alternative σ factor mediating response to nitrogen starvation (σ^54^ in *Escherichia coli*; σ^N^ in other bacteria; Yang et al., 2015; Glyde et al., 2018)--are members of the same protein family as group-1 σ factors (reviewed in Feklistov et al., 2014). Group-2 and group-3 alternative σ factors are closely related in structure to group-1 σ factors, lacking only functional modules σR1.1 (in group-2 σ factors) or σR1.1 and σR1.2 (in group-3 σ factors). The close structural similarity of group-2 and group-3 σ factors to group-1 σ factors, together with crystal structures of transcription initiation complexes containing group-2 σ factors (Liu et al., 2016), facilitates an understanding of the mechanism of group-2- and group-3-σ-factor-dependent transcription initiation.

Group-4 alternative σ factors--also referred to as “extracytoplasmic σ factors” (ECF σ factors), based on functional roles in response to cell-surface and other extracytoplasmic stresses--are only distantly related to group-1 σ factors and are substantially smaller than group-1 σ factors, lacking four of the six functional modules present in group-1 σ factors (Fig. 1A; reviewed in Lonetto et al., 1994; Missiakas and Raina, 1998; Helmann, 2002, 2016; Starori et al., 2009; Mascher, 2013; Feklistov et al., 2014; Campagne et al., 2015; Sineva and Ades, 2017). ECF σ factors comprise only a module related to σR2 (the module that recognizes promoter −10 elements in group-1 σ factors), a module related to σR4 (the module that recognizes promoter −35 elements in group-1 σ factors)., and a short σR2/4 linker that has no detectable sequence similarity to the σR3/4 linker of group-1 σ factors. No structural information previously has been reported for RNAP holoenzymes or transcription initiation complexes containing ECF σ factors. In the absence of structural information for ECF σ factors, it has been unclear how ECF σ factors, despite lacking sequences homologous to the σR3/4 linker of group-1 σ factors, are able to connect σR2 and σR4 with an appropriate spacing to recognize promoter −10 and −35 elements, are able to pre-organize the DNA template strand to facilitate initiating-nucleotide binding and *de novo* transcription initiation; and are able to coordinate entry of RNA into the RNA exit channel with promoter escape. In addition, in the absence of structural information, and with comparatively limited sequence similarity between σR2 of ECF σ factors and σR2 of group-1 σ factors (Feklistov et al., 2014), it has been unclear whether σR2 of ECF σ factors adopts the same fold as σR2 of group-1 σ factors and uses the same strategy to bind and unwind the promoter −10 element as group-1 σ factors.

ECF σ factors are numerically the largest, and functionally the most diverse, alternative σ factors (Missiakas and Raina, 1998; Helmann, 2002, 2016; Starori et al., 2009; Mascher, 2013; Feklistov et al., 2014; Campagne et al., 2015; Sineva and Ades, 2017). Fully ten of the thirteen σ factors in *Mycobacterium tuberculosis (Mtb),* the causative agent of tuberculosis, are ECF σ factors: σ^C^, σ^D^, σ^E^, σ^G^, σ^H^, σ^I^, σ^J^, σ^K^, σ^L^, and σ^M^, mediating responses to nutrition depletion, surface stress, temperature stress, oxidative stress, pH stress, growth in stationary phase, and growth in macrophages (Manganelli et al., 2004, 2013; Rodrigue et al., 2006; Sachdeva et al., 2010; Newton-Foot et al., 2013). For example, the *Mtb* ECF σ factor σ^L^ (Fig. S1A) mediates response to oxidative stress and regulates its own synthesis, polyketide-synthase synthesis, cell-wall synthesis, lipid transport, the oxidative state of exported proteins, and virulence (Hahn et al., 2005; Dainese et al., 2006; Rodrigue et al. 2007).

In this work, we have determined crystal structures, at 3.3 to 3.8 Å resolution of functional transcription initiation complexes comprising *Mtb* RNAP, the *Mtb* RNAP ECF σ factor σ^L^, and nucleic-acid scaffolds corresponding to the transcription bubble and downstream dsDNA of an ECF-σ-factor-dependent RNAP-promoter open complex *(Mtb* RPo-σ^L^), or an RNAP-promoter initial transcribing complex *(Mtb* RPitc-σ^L^) (Table 1; Figs. 1, S1–S2).

**Table 1.** Structure data collection and refinement statistics.

| structure | *Mtb* RPitc5-$\sigma^L$_sp4 | *Mtb* RPitc5-$\sigma^L$_sp5 | *Mtb* RPitc5-$\sigma^L$_sp6 |
| --- | --- | --- | --- |
| PDB code | 6DV9 | 6DVB | 6DVC |
| **data collection\*** |  |  |  |
| data collection | APS 19-ID | SSRL-9-2 | APS 19-ID |
| source |  |  |  |
| space group | P2<sub>1</sub>2<sub>1</sub>2<sub>1</sub> | P2<sub>1</sub>2<sub>1</sub>2<sub>1</sub> | P2<sub>1</sub>2<sub>1</sub>2<sub>1</sub> |
| cell dimensions |  |  |  |
| a, b, c (Å) | 143.3,161.4,237.7 | 143.7,160.6,240.4 | 146.3,161.5,240.6 |
| $\alpha$, $\beta$, $\gamma$ (°) | 90.0, 90.0, 90.0 | 90.0, 90.0, 90.0 | 90.0, 90.0, 90.0 |
| resolution (Å) | 50.0-3.8 (3.9-3.8) | 50.0-3.8 (3.9-3.8) | 50.0-3.3 (3.4-3.3) |
| number of unique reflections | 53,020 | 53,728 | 98,083 |
| R<sub>merge</sub> | 0.162 (0.567) | 0.188 (0.706) | 0.175 (0.710) |
| R<sub>meas</sub> | 0.172 (0.613) | 0.200 (0.752) | 0.184 (0.764) |
| R<sub>pim</sub> | 0.056 (0.222) | 0.066 (0.246) | 0.055 (0.273) |
| CC<sub>1/2</sub> (highest resolution shell) | 0.526 | 0.715 | 0.558 |
| I/σI | 8.6 (2.2) | 7.3 (1.9) | 13 (1.9) |
| completeness (%) | 96.6 (90.5) | 97.1 (96.5) | 98.9 (99.3) |
| redundancy | 10.7 (9.9) | 8.2 (8.1) | 10.4 (7.4) |
| anomalous | N/A | N/A | N/A |
| completeness(%) |  |  |  |
| anomalous | N/A | N/A | N/A |
| redundancy |  |  |  |
| **refinement\*** |  |  |  |
| resolution (Å) | 50.0-3.8 | 50.0-3.8 | 50.0-3.3 |
| number of unique reflections | 52,512 | 53,617 | 85,687 |
| number of test reflections | 2,625 | 2,691 | 4,267 |
| R<sub>work</sub>/R<sub>free</sub> | 0.18/0.23 (0.29/0.32) | 0.20/0.24 (0.32/0.35) | 0.19/0.23 (0.34/0.35) |
| number of atoms |  |  |  |
| protein | 24,956 | 24,974 | 24,982 |
| ligand/ion | 3 | 2 | 2 |
| r.m.s.deviation |  |  |  |
| bond lengths (Å) | 0.002 | 0.002 | 0.002 |
| bond angles (°) | 0.489 | 0.457 | 0.468 |
| MolProbity statistics |  |  |  |
| clash score | 7.2 | 6.3 | 6.0 |
| rotamer outliers (%) | 1.6 | 2.4 | 2.6 |
| Cβ outliers (%) | 0 | 0 | 0 |
| Ramachandran plot |  |  |  |
| favored (%) | 95.2 | 95.4 | 95.5 |
| outliers (%) | 0.3 | 0.2 | 0.3 |

\*Highest resolution shell in parentheses.

**Table 1. Structure data collection and refinement statistics (continued).**
| structure | *Mtb* [BrU]RPO-$\sigma^L$_sp6 | *Mtb* [SeMet15,76]RPO-$\sigma^L$_sp6 |
| --- | --- | --- |
| PDB code | 6DVD | 6DVE |
| **data collection\*** |  |  |
| data collection | APS 19-ID | APS 19-ID |
| source |  |  |
| space group | P2<sub>1</sub>2<sub>1</sub>2<sub>1</sub> | P2<sub>1</sub>2<sub>1</sub>2<sub>1</sub> |
| cell dimensions |  |  |
| a, b, c (Å) | 142.1, 161.5, 239.4 | 142.8, 160.6, 240.2 |
| $\alpha$, $\beta$, $\gamma$ (°) | 90.0, 90.0, 90.0 | 90.0, 90.0, 90.0 |
| resolution (Å) | 50.0-3.9 (4.0-3.9) | 50.0-3.8 (3.9-3.8) |
| number of unique reflections | 44,608 | 50,281 |
| R<sub>merge</sub> | 0.110 (0.768) | 0.178 (>1.000) |
| R<sub>meas</sub> | 0.121 (0.847) | 0.187 (>1.000) |
| R<sub>pim</sub> | 0.047 (0.348) | 0.062 (0.608) |
| CC<sub>1/2</sub> (highest resolution shell) | 0.797 | 0.589 |
| I/σI | 14.3 (1.6) | 10.2 (1.0) |
| completeness (%) | 87.4 (68.5) | 92.1 (79.3) |
| redundancy | 6.0 (4.8) | 9.3 (5.5) |
| anomalous | 87.4 | 92.1 |
| completeness(%) |  |  |
| anomalous | 6.0 | 9.3 |
| redundancy |  |  |
| **refinement\*** |  |  |
| resolution (Å) | 45.9-3.9 | 46.7-3.8 |
| number of unique reflections | 37,593 | 40,877 |
| number of test reflections | 1,996 | 1,987 |
| R<sub>work</sub>/R<sub>free</sub> | 0.22/0.24 (0.25-0.26) | 0.20/0.24 (0.25/0.30) |
| number of atoms |  |  |
| protein | 24,743 | 24,782 |
| ligand/ion | 2 | 4 |
| r.m.s.deviation |  |  |
| bond lengths (Å) | 0.002 | 0.002 |
| bond angles (°) | 0.491 | 0.492 |
| MolProbity statistics |  |  |
| clash score | 6.6 | 6.8 |
| rotamer outliers (%) | 2.6 | 1.7 |
| Cβ outliers (%) | 0 | 0 |
| Ramachandran plot |  |  |
| favored (%) | 95.3 | 95.0 |
| outliers (%) | 0.3 | 0.3 |

\*Highest resolution shell in parentheses.

## RESULTS

### Structures of *Mtb* RPo-σ^L^ and *Mtb* RPitc-σ^L^

Structures were determined using recombinant *Mtb* RNAP core enzyme prepared by co-expression of *Mtb* RNAP subunit genes in *E. coli,* recombinant *Mtb* σ^L^, and synthetic nucleic-acid scaffolds based on the sequence of the σ^L^-dependent promoter *P-sigL* (the promoter responsible for expression of the gene encoding σ^L^; Hahn et al., 2005; Dainese et al., 2006; Rodrigue et al. 2007) (Figs. S1–S2). Transcription experiments demonstrate that *Mtb* RNAP-σ^A^ holoenzyme (containing the group-1 σ factor σ^A^) does not efficiently perform transcription initiation at the P*-sigL* promoter, whereas *Mtb* RNAP-σ^L^ holoenzyme (containing the ECF σ factor σ^L^) does (Fig. S1E). We prepared “downstream-fork-junction” nucleic-acid scaffolds containing P-sigL sequences, analogous to the downstream-fork-junction nucleic-acid scaffolds containing consensus group-1-σ-factor-dependent promoter sequences used previously for structural analysis of group-1-σ-factor-dependent transcription initiation (Fig. S2, left panels). Because the P-sigL transcription start site (TSS) had been mapped only provisionally (Hahn et al., 2005; Dainese et al., 2006; Rodrigue et al., 2007), we prepared and analyzed a set of downstream-fork-junction nucleic-acid scaffolds having different lengths--4 nt, 5 nt, 6, or 7 nt--of the “spacer” between the P-sigL promoter −10 region and downstream dsDNA (Fig. S2, left panels). Transcription experiments indicated that all analyzed nucleic-acid scaffolds were functional in σ^L^-dependent *de novo* transcription initiation at the expected TSS (with the initiating nucleotide base-pairing to template-strand ssDNA 2 nt upstream of dsDNA), and σ^L^-dependent primer-dependent transcription initiation at the expected TSS (with the primer 3’ nucleotide base-pairing to template-strand ssDNA 2 nt upstream of dsDNA), with highest levels of function observed for a spacer length of 6 nt (Fig. S1F-G). Robotic crystallization trials identified crystallization conditions yielding high-quality crystals for spacer lengths of 4 nt, 5 nt, or 6 nt (Table 1; Fig. S2, center panels). X-ray datasets were collected at synchrotron beam sources, and structures were solved by molecular replacement and refined to 3.3 to 3.8 Å resolution (Table 1; Fig. S2, right panels).

Experimental electron-density maps showed clear density for RNAP, σ^L^, and nucleic acids (Fig. S2, right panels). The resulting structures were essentially identical for nucleic-acid scaffolds having spacer lengths of 4 nt, 5 nt, or 6 nt (Figs. S2, right panels). However, map quality was highest for the nucleic-acid scaffold having a spacer length of 6 nt, and, therefore subsequent analysis focussed on structures with a spacer length of 6 nt *(Mtb* RNAP-σ^L^ RPitc5_sp6). For the nucleic-acid scaffold containing a 6 nt spacer, the translocational state of the transcription complex was experimentally verified by preparation of a scaffold having a single 5-bromo-dU substitution and collection of bromine anomalous diffraction data (Table 1; Fig. S2D). The fit of σ^L^ separately was experimentally verified by preparation of a selenomethionine-labelled σ^L^ derivative and collection of selenium anomalous diffraction data (Table 1; Fig. S2E).

### Protein-protein interactions between ECF σ factor and RNAP

The structural organization of the ECF σ^L^-factor-dependent transcription initiation complex is unexpectedly similar to that of a group-1 σ^A^-factor-dependent transcription initiation complex (Figs. 1B, 2). σR2 and σR4 of σ^L^ occupy the same positions on RNAP, and make the same interactions with RNAP, as σR2 and σR4 of σ^A^ factor (Fig. 2). Despite the much smaller size of the connector between σR2 and σR4 in σ^L^ as compared to σ^A^ (20 residues vs. 84 residues; Fig. S1A), the connector in σ^L^ spans the full distance between the σR2 and σR4 binding positions on RNAP and follows a path through RNAP remarkably similar to that of the connector in σ^A^ (Fig 2.). Thus, the σ^L^ σR2/4 linker, like the σ^A^ σR3/4 linker, first enters the RNAP active-center cleft and approaches the RNAP active center, and then makes a sharp turn and exits the RNAP active-center cleft through the RNAP RNA-exit channel.

**Figure 2.**
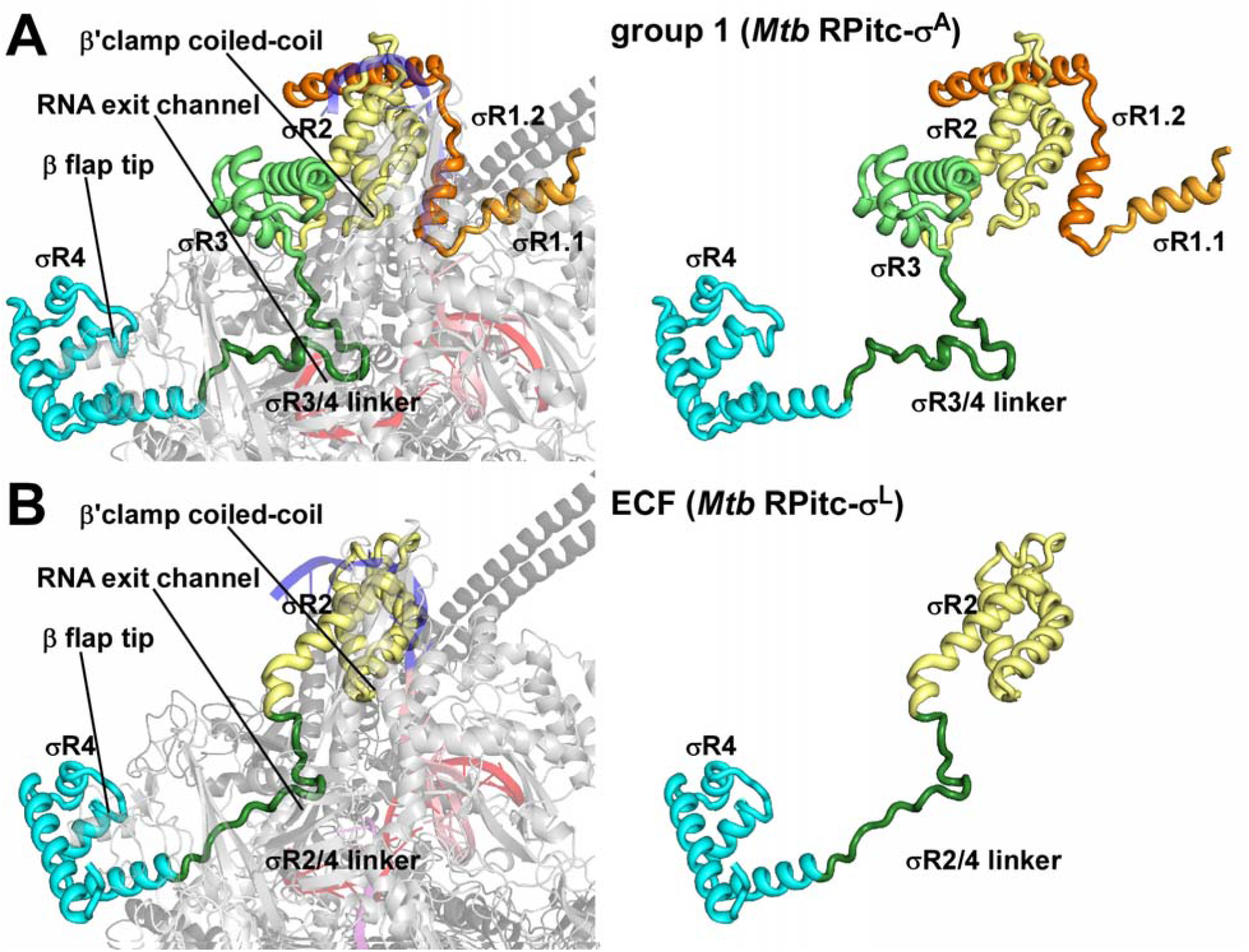
Comparison of protein-protein interactions by group-1 and ECF σ factors with RNAP core enzyme. **(A)** Protein-protein interactions by group 1 (σ^A^) σ factor. **(B)** Protein-protein interactions by ECF (σ^L^) σ factor. Colors are as in Fig. 1. See Fig. S7.

Inside the RNAP active-center cleft, the σ^L^ σR2/4 linker, like the σ^A^ σR3/4 linker, makes direct interactions with template-strand ssDNA nucleotides of the unwound transcription bubble (Figs. 2–4, S3A). The interactions of the σ^L^ σR2/4 linker with template-strand ssDNA include a direct H-bonded interaction of σ^L^ Ser96 with a Watson-Crick H-bonding atom of the template-strand nucleotide at promoter position −5 (Figs. 3B and S3A, bottom). The interactions of the σ^L^ σR2/4 linker with template-strand ssDNA are similar to, but less extensive than, those of the σ^A^ σR3/4 linker with template-strand ssDNA, which include direct H-bonded interactions of σ^A^ Asp432 and Ser433 with Watson-Crick H-bonding atoms of template-strand ssDNA nucleotides at promoter positions −4 and −3 (Figs. 3A and S3A).

**Figure 3.**
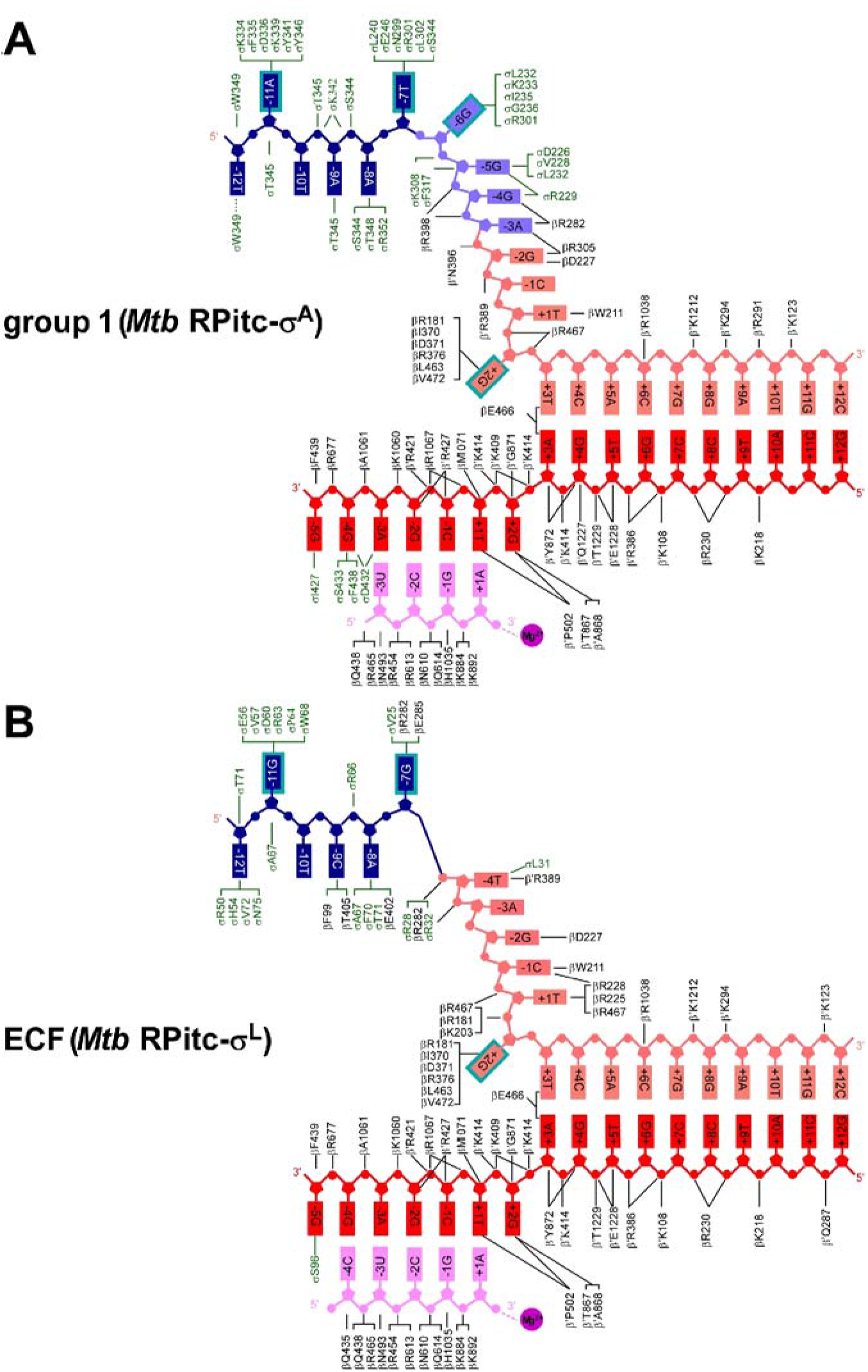
Comparison of protein-nucleic acid interactions with group-1 and ECF σ factors: summary. **(A)** Summary of protein-nucleic acid interactions in *Mtb* RPitc-σ^A^. Black residue numbers and lines, interactions by *Mtb* RNAP; green residue numbers and lines, interactions by *Mtb* σ^A^; blue, −10 element of DNA nontemplate strand; light blue, discriminator element of DNA nontemplate strand; pink, rest of DNA nontemplate strand; red, DNA template strand; magenta, RNA product; violet circle, RNAP active-center Mg^2+^; cyan boxes, bases unstacked and inserted into protein pockets. Residues are numbered as in *Mtb* RNAP. **(B)** Summary of protein-nucleic acid interactions in *Mtb* RPitc-σ^L^. Green residue numbers and lines, interactions by *Mtb* σ^L^. Other colors are as in (A). See Figs. S3–S5 and S7.

**Figure 4.**
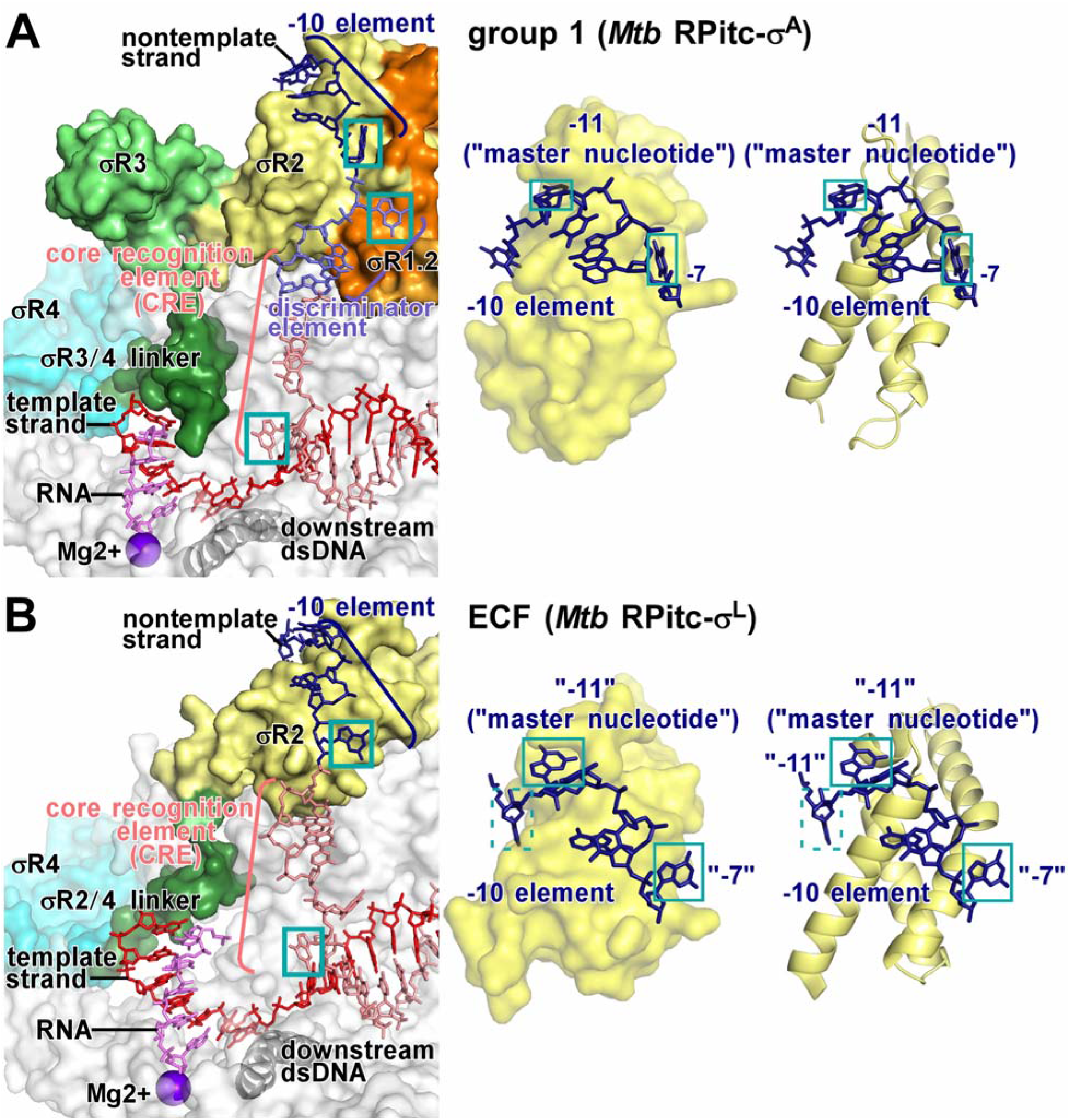
Comparison of protein-nucleic acid interactions with group-1 and ECF σ factors: interactions with transcription-bubble nontemplate and template strands. **(A)** Left: interactions of *Mtb* RNAP and σ^A^ with transcription-bubble nontemplate strand, transcription-bubble template strand, and downstream dsDNA. Right: interactions of σ^A^ σR2 with σ^A^-dependent promoter −10 element. For promoter positions −11 (“master nucleotide”; Lim et al., 2001) and −7, bases are unstacked and inserted into pockets (cyan boxes). Colors are as in Figs. 1–3. **(B)** Left: interactions of *Mtb* RNAP and σ^L^ with transcription-bubble nontemplate strand, transcription-bubble template strand, and downstream dsDNA. Right: interactions of σ^L^ σR2 with σ^L^-dependent promoter −10 element. For two promoter positions, here designated “−11” (“master nucleotide”) and “−7,” by analogy to corresponding nucleotides in group-1-σ-factor complex (panel A), bases are unstacked and inserted into pockets (cyan boxes). For one additional nucleotide, here designated “−12,” the base also appears to be unstacked and inserted into a pocket (dashed cyan box). See Figs. S4–S5 and S7.

In the case of the group-1 σ factor, σ^A^, the interactions between this segment of the σR3/4 linker and template-strand ssDNA pre-organize template-strand ssDNA to adopt a helical conformation and to engage the RNAP active-center nucleotide-addition site, thereby facilitating initiating-nucleotide binding and *de novo* initiation (Zhang et al., 2012; see also Kulbachinskiy and Mustaev, 2006; Pupov et al., 2014). The similarity of the interactions made by the ECF σ factor, σ^L^, suggests that ECF σ factors likewise pre-organize template-strand ssDNA and facilitate initiating-nucleotide binding and *de novo* initiation.

In the case of the group-1 σ factor, σ^A^, the interactions between this segment of the σR3/4 linker and template-strand ssDNA must be broken, and this segment of the σR3/4 linker must be displaced, when the nascent RNA reaches a length >4 nt during initial transcription, and this requirement for breakage of interactions and displacement is thought to impose an energy barrier that results in, or enhances, abortive initiation (Murakami et al., 2002; Kulbachinskiy and Mustaev, 2006; Zhang et al., 2012; Basu et al., 2014; Pupov et al., 2014) and initial-transcription pausing (Duchi et al., 2016; Lerner et al., 2016; Dulin et al., 2018). The similarity of the interactions made by the ECF σ factor, σ^L^, suggests that ECF σ factors likewise have a similar requirement for displacement of a linker segment during initial transcription--in this case, when the nascent RNA reaches a length of >5 nt (Fig. S3B)--and that this similar requirement imposes a energy barrier that results in, or enhances, abortive initiation and initial-transcription pausing. Consistent with this hypothesis, transcription and transcript-release experiments indicate that *Mtb* RNAP-σ^L^ holoenzyme efficiently performs abortive initiation, efficiently producing and releasing short abortive RNA products (Fig. S3C).

The 5 C-terminal residues of the σ^L^ σR2/4 linker, like the 10 C-terminal residues of the σ^A^ σR3/4 linker, exit the RNAP active-center cleft and connect to σR4 by threading through the RNAP RNA-exit channel (Fig. 2). In the case of the group-1 σ factor, σ^A^, the C-terminal segment of the σR3/4 linker must be displaced from the RNA-exit channel when the nascent RNA reaches a length of 11 nt at the end of initial transcription and moves into the RNA-exit channel, and this displacement is thought to alter interactions between σR4 and RNAP, and thereby to trigger promoter escape and to transform the transcription initiation complex into a transcription elongation complex (Murakami et al., 2002; Vassylyev et al, 2002; Mekler et al., 2002). The similarity of the threading through the RNAP RNA-exit channel by the ECF σ factor, σ^L^, suggests that ECF σ factors have a similar requirement for displacement of a linker C-terminal segment and have a similar mechanism of promoter escape and transformation from transcription initiation complexes into transcription elongation complexes.

In its interactions with template-strand ssDNA and the RNAP RNA exit channel, the σ^L^ σR2/4 linker, like the σ^A^ σR3/4 linker, linker serves as a molecular mimic, or a molecular placeholder, for nascent RNA, making interactions with template-strand ssDNA and the RNAP-RNA exit channel in early stages of transcription initiation that subsequently, in late stages of transcription initiation and in transcription elongation, are made by nascent RNA. The σ^L^ σR2/4 linker and the σ^A^ σR3/4 linker, both have net negative charge (Fig. S1A), and both employ extended conformations (fully extended for the σ^L^ σR2/4 linker; largely extended for the σ^A^ σR3/4 linker; Fig. 2) to interact with template-strand ssDNA and the RNAP RNA exit channel, consistent with function as molecular mimics of a negatively charged, extended nascent RNA. Nevertheless, the σ^L^ σR2/4 linker and the σ^A^ σR3/4 linker exhibit no detectable sequence similarity (Fig. S1A) and no detailed structural similarity (Fig. 2). We conclude that the σR2/4 linker of an ECF σ factor and the σR3/4 linker of a group-1 σ factor provide an example of functional analogy in the absence of structural homology.

### Protein-DNA interactions between ECF σ factor and promoter: −10 element

The structure reveals the interactions between the ECF σ factor, σ^L^, and promoter DNA that mediate recognition of the promoter −10 element (Figs. 3–5, S4). The σ^L^ conserved module σR2, like the σ^A^ conserved module σR2, mediates recognition of the promoter −10 element through interactions with nontemplate-strand ssDNA in the unwound transcription bubble (Figs. 3–4). In the case of the group-1 σ factor, σ^A^, a crucial aspect of recognition of the promoter −10 element is unstacking of nucleotides, flipping of nucleotides, and insertion of nucleotides into protein pockets at two positions of the σ^A^-dependent promoter: i.e., position −11 (referred to as the “master nucleotide,” based on its especially important role in promoter recognition; Lim et al., 2001) and position −7 (Figs. 3A, 4A, S4). The ECF σ factor, σ^L^, unstacks nucleotides, flips nucleotides, and inserts nucleotides into protein pockets at the corresponding positions of the σ^L-^dependent promoter (here designated positions “−11” and “−7”; Figs. 3B, 4B, 5, S4) and also unstacks and inserts a nucleotide into a protein pocket at one additional position of the σ^L^-dependent promoter (position “−12”; Figs. 4B, 5).

**Figure 5.**
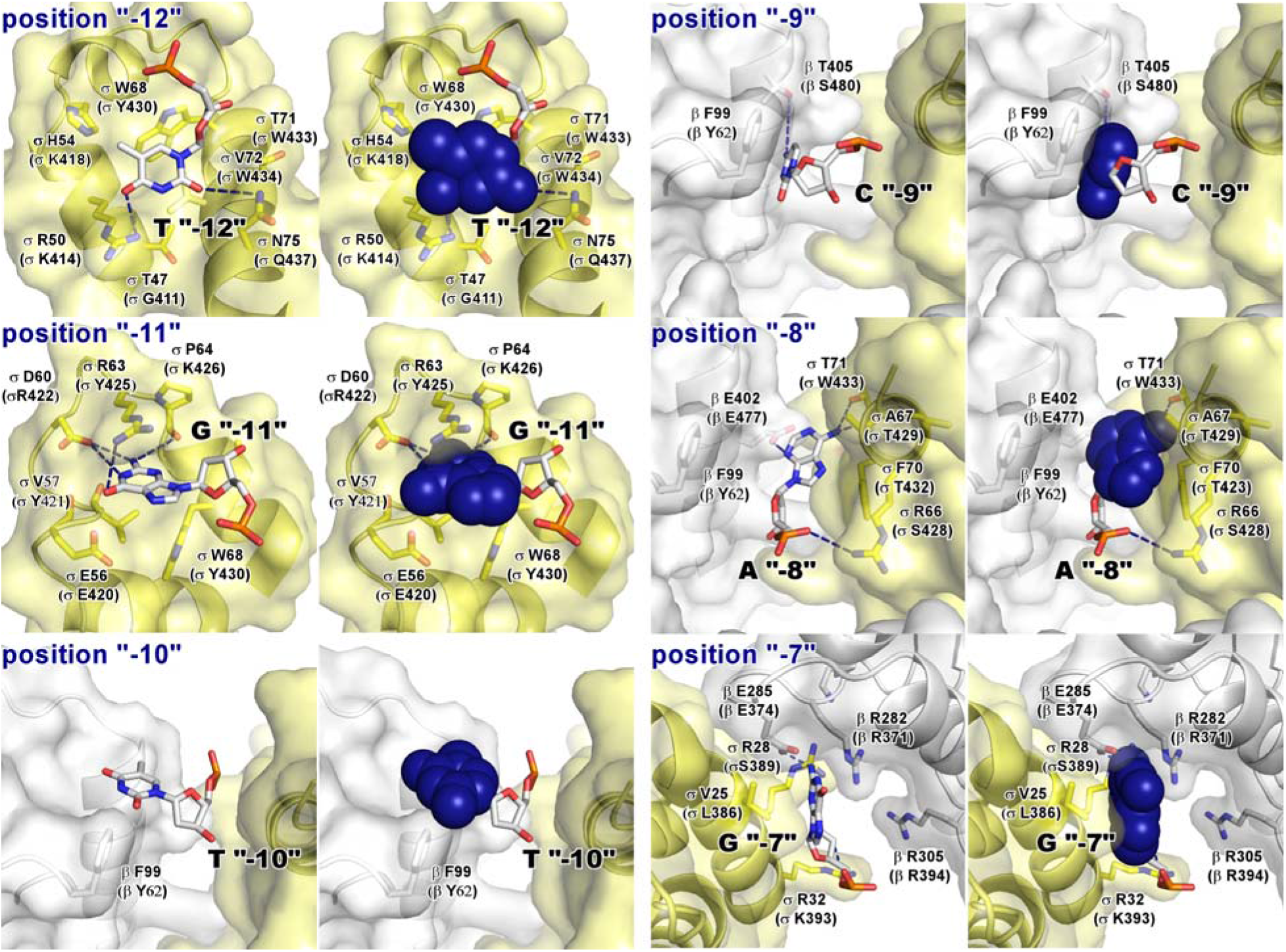
Recognition by *Mtb* σ^L^ of σ^L^-promoter −10 element: interactions with nontemplate strand positions “−12” through “−7”. For each promoter position, left subpanel shows DNA nucleotides in stick representation to highlight individual protein-nucleotide interactions, and right panel shows DNA nucleotide base moieties in space-filling representation to highlight protein-base steric complementarity. Yellow surfaces, solvent-accessible surfaces of *Mtb* σ^L^; gray surfaces, solvent-accessible surfaces of *Mtb* RNAP β subunit; dark blue surfaces, van der Waals surfaces of σ^L^-promoter −10 element; yellow ribbons, *Mtb* σ^L^ backbone; gray ribbons, *Mtb* RNAP β subunit backbone; yellow, yellow-blue, and yellow-red stick representations, σ^L^ carbon, nitrogen, and oxygen atoms, respectively; white, blue, red, and orange stick representations, DNA carbon, nitrogen, oxygen, and phosphorous atoms, respectively; blue dashed lines, H-bonds. Residues are numbered as in *Mtb* RNAP and σ^L^, and, in parentheses, as in *E. coli* RNAP and σ^70^. See Fig. S4.

RNAP σ^L^ holoenzyme unstacks, flips, and inserts into a protein pocket a guanosine at position “−11” of the σ^L^-dependent promoter, making extensive interactions with the base moiety of the guanosine, including multiple direct H-bonded interactions with Watson-Crick H-bonding atoms (Figs. 3, 4B, 5, S4A). The interactions between σ^L^ and guanosine at position “−11” of the σ^L^-dependent promoter are similar to the interactions between RNAP σ^A^ holoenzyme and adenosine at position −11 of the σ^A^-dependent promoter −10 element, including, in particular, similar stacking interactions of σ^L^ aromatic amino acid Trp48 with guanosine and of corresponding σ^A^ aromatic amino acid Tyr436 with adenosine (Fig. S4A). The different specificities--guanosine at position “−11” for σ^L^ vs. adenosine at position −11 for σ^A^--arise from differences in H-bond-donor/H-bond-acceptor character of atoms forming the floors of the relevant protein pockets of σ^L^ and σ^A^, with H-bonding complementarity to guanosine in σ^L^ and H-bonding complementarity to adenosine in σ^A^ (Fig. S4A).

RNAP σ^L^ holoenzyme also unstacks, flips, and inserts into a pocket a guanosine at position “−7” of the σ^L^-dependent promoter, making extensive interactions with the base moiety of the guanosine, including a direct H-bonded interaction with a Watson-Crick H-bonding atom (Figs. 3, 4B, 5, S4B).

These interactions are similar in location to, but different in detail from the interactions made by RNAP σ^A^ holoenzyme with thymidine at position −7 of the σ^L^-dependent promoter (Figs. 4, S4B). The differences in detail arise from the fact that σ^L^ does not contain conserved module σR1.2. In the case of σ^L^, the interactions involve residues of σR2 and residues of RNAP β subunit, with the base moiety of the guanosine at position “−7” being inserted into a cleft between σR2 and β (Figs. 5, S4B). In contrast, in the case of σ^A^, the interactions involve residues of σR2 and residues σR1.2, with the base moiety of the thymidine at position −7 being inserted into a cleft between σR2 and σR1.2 (Fig S4B).

RNAP σ^L^ holoenzyme also appears to unstack and insert into a protein pocket a thymidine at position “−12” of the σ^L^-dependent promoter (Fig. 4B, 5), placing one face of the base moiety of the thymidine in a shallow surface pocket, in position to make a direct H-bonded interaction with a Watson-Crick atom (Fig. 5). The interaction with an unstacked nucleotide inserted into a protein pocket implies that position “−12” of the σ^L^-dependent promoter must be ssDNA in RPo-σ^L^ and RPitc-σ^L^, and thus that the transcription bubble must extend to position “−12” in RPo-σ^L^ and RPitc-σ^L^. This interaction does not have a counterpart in the σ^A^-dependent transcription initiation complex, in which position −12 of the σ^A^-dependent promoter is dsDNA and in which the transcription bubble extends only to position −11 (Bae et al., 2015; Zuo and Steitz, 2015; Feng et al, 2016).

In addition to these potential specificity-determining interactions with unstacked nucleotides inserted into protein pockets, RNAP σ^L^ holoenzyme makes potentially specificity-determining interactions with positions “−9” and “−8” of the σ^L^-dependent promoter (Fig. 5). RNAP σ^L^ holoenzyme makes a direct H-bonded interaction, through RNAP β subunit, with a Watson-Crick atom of the base moiety of cytidine at position “−9” (Fig. 5) and makes two direct H-bonded interactions, through σR2 and RNAP β subunit, with a Watson-Crick atom of the base moiety of adenosine at position “−8” (Fig. 5).

Biochemical experiments assessing effects of all possible single-base-pair substitutions at each position of the *P-sigL* promoter −10 region confirm the functional significance of the positions contacted in the crystal structure (i.e., positions“−12,” “−11,” “−9,” “−8,” and “−7”; Fig. 6A), confirm the sequence preferences at these positions inferred from the crystal structure (Fig. 6A), and yield a revised consensus sequence for the σ^L^-dependent −10 element of T_“−12”_-G_“−11”-_N_“−10”-_C/A_“−9”-_A_“−8”-_G_“−7”_ (Fig. 6B). The revised consensus sequence for the σ^L^-dependent −10 element matches the literature consensus sequence (Hahn et al, 2005; Sachdeva et al., 2009; Manganelli, 2013; Newton-Foot, 2013), in its first four positions (T_“−12”-_G_“−11”-_N_“−10”_-C/A_“−9”_) and extends the literature consensus sequence for two additional positions (A_“−8”-_G_“−7”_). Consistent with the observed extensive network of H-bonded interactions involving position “−11G” (Figs. 3, 5, S4A), specificity is observed to be strongest at position “−11” (Figs. 6A-B).

**Figure 6.**
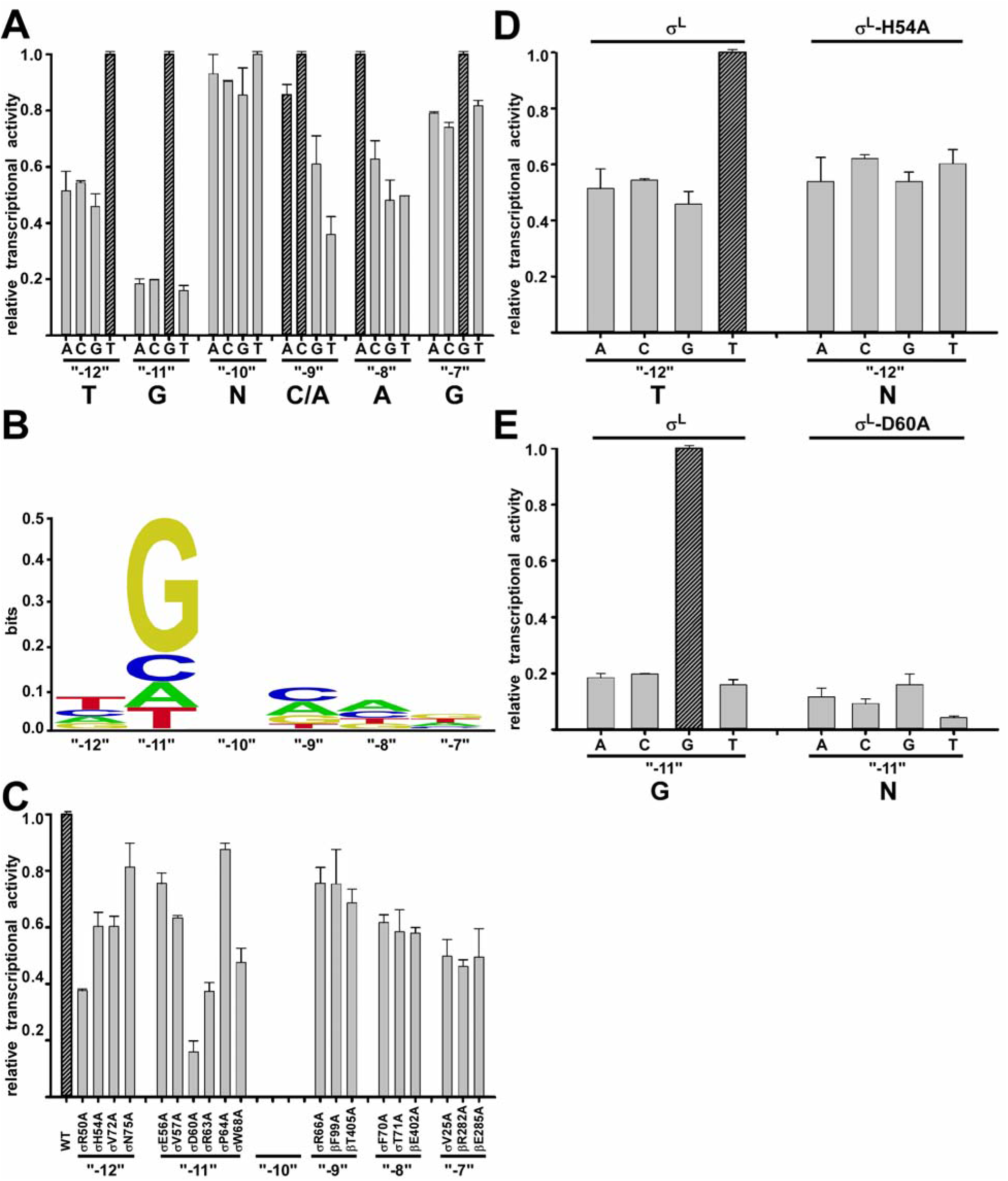
Recognition by *Mtb* σ^L^ of σ^L^-promoter-10 element: experimental data. **(A)** Systematic-substitution experiments defining σ^L^-dependent promoter-10-element consensus sequence. Relative transcription activities of derivatives of σ^L^-dependent promoter P-*sigZ* having all possible single base-pair substitutions at each position of promoter −10 element,”-12” through “ −7.” Inferred consensus nucleotides are shown at bottom, and data for inferred consensus nucleotides are hatched. **(B)** Sequence logo for σ^L^-promoter −10-element consensus sequence [generated using transcription data from (A) and enoLOGOS (Workman et al., 2005; http://biodev.hgen.pitt.edu/enologos/; input setting “energy (2)” and weight-type setting “probabilities.”] **(C)** Alanine-scanning experiments (Cunningham and Wells, 1989) demonstrating functional importance of observed amino acid-base interactions in recognition of σ^L^-promoter −10 element. Effects on transcription of alanine substitutions of σ^L^ amino acids that contact σ^L^-dependent promoter −10 element, positions “−12” through “−7” (identities of contacting amino acids from Figs. 3 and 5). **(D)-(E)** Loss-of-contact experiments (Ebright, 1985, 1986, 1991; Zhang et al., 1990) indicating that σ^L^ residues His54 and Asp60 determine specificity at position “−12” and “−11,” respectively. Left: transcriptional activity with wild-type σ^L^ for all possible single base-pair-substitutions at indicated position (strong specificity for consensus base pair). Right: transcriptional activity of σ^L^ derivatives having alanine substitutions (no specificity for consensus base pair). See Fig. S6.

“Alanine-scanning” experiments (Cunningham and Wells, 1989), in which residues of σ^L^ that contact −10-element nucleotides in the crystal structure are substituted with alanine and effects on σ^L^-dependent transcription are quantified, confirm the functional significance of the observed interactions (Fig. 6C).

“Loss-of-contact” experiments (Ebright, 1985, 1986, 1991; Zhang et al., 1990), in which residues of σ^L^ that contact −10-element nucleotides in the crystal structure are substituted with alanine and effects on specificity at the contacted positions are quantified, confirm that σ^L^ His54 determines specificity for thymidine at position “−12” (Fig. 6D) and that σ^L^ Asp60 determines specificity for guanosine at position “−11” (Fig. 6E). In the crystal structure, σ^L^ His54 makes a van der Waals interaction with the 5’-methyl group of the base moiety of thymidine at position “−12” (Fig. 5); in loss-of-contact experiments, substitution of His54 by alanine eliminates specificity for thymidine at position “−12” (Fig. 6D). In the crystal structure σ^L^ Asp60 makes an H-bonded interaction with Watson-Crick atoms of the base moiety of guanosine G at position “−11” (Figs. 5, S4A; in loss-of-contact experiments, substitution of Asp60 by alanine eliminates specificity for thymidine at position “−11” (Fig. 6E).

### Protein-DNA interactions between ECF σ factor and promoter: core recognition element (CRE)

The structure reveals the interactions between RNAP σ^L^ holoenzyme and nontemplate-strand ssDNA downstream of the promoter −10 element in the ECF σ^L^-dependent transcription initiation complex (Figs. 3B, S5). In the case of group-1-σ-factor-dependent transcription initiation complexes, sequence-specific interactions occur between RNAP β subunit and a 6 nt segment of nontemplate-strand ssDNA downstream of the promoter −10 element referred to as the “core recognition element” (CRE; positions −6 through +2; Zhang et al., 2012; Lin et al., 2017; Vahedian-Movahed, 2017). These interactions include, most notably, (1) stacking of a tryptophan residue of RNAP β subunit on the base moiety of thymidine at nontemplate-strand position +1 (Fig. S5A), and (2) unstacking, flipping, and insertion into a protein pocket, formed by the RNAP β subunit, of the guanosine at nontemplate-strand position +2 (Figs. 3, 4A, S5B). The identical interactions occur in the ECF σ^L^-dependent transcription initiation complex (Figs. 3, 4B, S5).

Biochemical experiments assessing effects of all possible base-pair substitutions at positions downstream of the P-sigL promoter −10 element (positions −4 through +2) confirm the functional significance of the interactions in the crystal structure with thymidine at position +1 and guanosine at position +2 (Fig. S6A) and yield a CRE consensus sequence for a ECF σ^L^-dependent transcription initiation complex (Fig. S6B) similar to the CRE consensus sequence for a group-1-σ-factor-dependent transcription initiation complex (Zhang et al., 2012; Vahedian-Movahed, 2017).

## DISCUSSION

Our structural results show that: (1) σR2 and σR4 of an ECF σ factor σ^L^ adopt the same folds and interact with the same sites on RNAP as σR2 and σR4 of a group-1 σ factor (Figs. 1–2); (2) the connector between σR2 and σR4 of ECF σ factor σ^L^ enters the RNAP active-center cleft to interact with template-strand ssDNA and then exits the RNAP active-center cleft by threading through the RNAP RNA-exit channel in a manner functionally analogous--but not structurally homologous--to the connector between σR2 and σR4 of a group-1 σ factor (Figs. 1–2; S3); (3) ECF σ factor σ^L^ recognizes the −10 element of an σ^L^-dependent promoter by unstacking nucleotides and inserting nucleotides into protein pockets at three positions of the transcription-bubble nontemplate-strand ssDNA (positions “−12,” “−11,” and “−7”; Figs. 3–5, S4), and (4) RNAP recognizes the core recognition element (CRE) of an σ^L^-dependent promoter by stacking a nucleotide on a tryptophan and by unstacking, flipping, and inserting a nucleotide into a protein pocket (positions +1 and +2; Figs. 3–4, S5). Our biochemical results confirm the functional significance of the observed protein-DNA interactions with the −10 element and CRE of an σ^L^-dependent promoter (Figs. 6A, S6A), provide consensus sequences for the −10 element and CRE of an σ^L^-dependent promoter (Figs. 6B, S6B), and define individual specificity-determining amino-acid-base interactions for two positions of the −10 element of an σ^L^-dependent promoter (positions “−12” and “−11”; Fig. 6C-D). The results provide an indispensable foundation for understanding the structural and mechanistic basis of ECF-σ-factor-dependent transcription initiation.

Our results regarding the connector between σR2 and σR4 of an ECF σ factor, in conjunction with previous results, indicate that all classes of bacterial σ factors contain structural modules that enter the RNAP active-center cleft to interact with template-strand ssDNA and then leave the RNAP active-center cleft by threading through the RNAP RNA-exit channel, providing mechanisms to facilitate *de novo* initiation, to coordinate extension of the nascent RNA with abortive initiation and initial-transcription pausing, and to coordinate entry of RNA into RNA-exit channel with promoter escape. For ECF σ factors, as shown here, the relevant structural module is the σR2/4 linker (Figs 3–4, S3); for group-1, group-2, and group-3 σ factors, the module is the functionally analogous--but not structurally homologous--σR3/4 linker (Murakami et al., 2002, 2013; Vassylyev et al, 2002; Zhang et al., 2012, 2014; Bae et al., 2013, 2015; Basu et al., 2014; Zuo and Steitz, 2015; Liu et al., 2016); and for group-σ^54^/σ^N^ σ factors, the module is the functionally analogous--but not structurally homologous--region II.3 (RII.3; Yang et al., 2015; Glyde et al., 2018).

More broadly, our results, in conjunction with previous results, indicate that cellular transcription initiation complexes in *all* organisms--bacteria, archaea, and eukaryotes--contain structural modules that enter the RNAP active-center cleft to interact with template-strand ssDNA and then leave the RNAP active-center cleft by threading through the RNAP RNA exit channel. In different classes of bacterial transcription initiation complexes, as described in the preceding paragraph, these roles are performed by the functionally analogous--but not structurally homologous--σR2/4 linker, σR3/4 linker, and RII.3. In archaeal transcription initiation complexes, these roles are performed by the TFIIB zinc ribbon and CSB, which are unrelated to the σR2/4 linker, σR3/4 linker, and RII.3 (Renfrow et al., 2004). In eukaryotic RNAP-I-, RNAP-II-, and RNAP-III-dependent transcription initiation complexes, these roles are performed by the Rm7 zinc ribbon and B-reader, the TFIIB zinc ribbon and B-reader, and the Brf1 zinc ribbon, respectively, each of which is unrelated to the σR2/4 linker, σR3/4 linker; and RII.3 (Kostrewa et al., 2009; Liu et al., 2010; He et al., 2016; Ptaschka et al., 2016; Engel et al., 2017; Han et al., 2017; Sadian et al., 2017; Vorländer et al., 2018; Abascal-Palacios et al., 2018). It is extraordinary that non-homologous, structurally and phylogenetically unrelated, structural modules are used to perform the same roles in different transcription initiation complexes, and is unknown how or why this occurs.

Our results define the protein-DNA interactions that ECF σ factor σ^L^ uses to recognize the −10 element of a σ^L^-dependent promoter. The consensus sequence obtained in this work for the −10-element of a σ^L^-dependent promoter, T_“−12”-_G_“−11”-_N_“−10”-_C/A_“−9”-_A_“−8”-_G_“−7”_ (Fig. 6B), confirms and extends the literature-consensus sequence (Hahn et al, 2005; Sachdeva et al., 2009; Manganelli, 2013; Newton-Foot, 2013), and the structural data of this work account for specificity at each specified position of the consensus sequence (Figs. 3–5, S4).

Previous work indicates that RNAP-σ^L^ holoenzyme prefers a C-G sequence immediately upstream of the −10-element (C_“--14”-_G_“−13”_ in our numbering system; Hahn et al, 2005; Sachdeva et al., 2009; Manganelli, 2013; Newton-Foot, 2013). Further previous work indicates that this preference may be shared by many RNAP-ECF-σ-factor holoenzymes (Helmann, 2002, 2016; Rodrigue et al., 2006, 2007; Sachdeva et al., 2009; Manganelli, 2013; Newton-Foot, 2013; Rhodius et al., 2013); for example, at least 8 of 10 *Mtb* RNAP-ECF-σ-factor holoenzymes--Mtb RNAP-σ^C^, −σ^D^, −σ^E^, −σ^G^, −σ^H^, −σ^J^, −σ^L^, and −σ^M^ holoenzymes--exhibit this preference (Rodrigue et al., 2006, 2007; Sachdeva et al., 2009; Manganelli, 2013; Newton-Foot, 2013). In this work, we performed crystallization using nucleic-acid scaffolds that did not contain C_“−14”-_G_“−13”_, and therefore our crystal structures do not definitively account for the preference for C_“−14”-_G_“−13”_. However, with the assumption that template-strand nucleotides at positions “^-^14” and “−13” of a σ^L^-dependent transcription initiation complex are positions similar to those in a group-1-σ-factor-dependent transcription initiation complex (Zuo and Steitz, 2015; Bae et al., 2015; Feng et al., 2016), our crystal structures suggest that the C-terminal α-helix of σ^L^ σR2 (σ^L^ residues 78-82) potentially could make direct, specificity-determining contacts with template-strand nucleotides at these positions. A similar mechanism for recognition of C-G immediately upstream of the −10 element has been proposed for the group-3 σ factor *E. coli* σ^28^ (Koo et al., 2009).

Both our structural results and our biochemical results point to the special importance of the nontemplate-strand nucleotide at position “−11” (“master nucleotide”; Figs. 3–6, S4A). Our results regarding recognition of the “−11” “master nucleotide” by an ECF σ factor are consistent with the NMR structure of a complex comprising σR2 from the *E. coli* ECF σ factor σ^E^ and a 5 nt oligodeoxyribonucleotide corresponding to part of the nontemplate strand of the −10 element of a σ^E^-dependent promoter (Campagne et al., 2014, 2015). The NMR structure showed unstacking, flipping, and insertion into a protein pocket of the “−11” “master-nucleotide” (a cytidine, rather than a guanosine, reflecting the different specificities of *E. coli* σ^E^ vs. *Mtb* σ^L^; Campagne et al., 2014, 2015). The NMR structure did not show unstacking and flipping of the nucleotide at position “−7,” reflecting the fact that the oligodeoxyribonucleotide in the NMR structure did not extend to position “−7” (Campagne et al., 2014, 2015). The NMR structure also did not show unstacking of the nucleotide at position “−12,” (Campagne et al., 2014, 2015), possibly reflecting an uncertainty in the NMR structure, or possibly reflecting a difference between *E. coli* σ^E^ and *Mtb* σ^L^ in recognition of position “−12.”

Based on the NMR structure, Campagne et al. (2014, 2015) hypothesized that the loop of σR2 that forms the protein pocket into which the “−11” “master nucleotide” is inserted--”loop L3” (residues 63-72 of *E. coli* σ^E^, which correspond to residues 56-67 of *Mtb* σ^L^)--serves as a functionally independent, functionally modular, determinant of specificity at the “master-nucleotide” position, such that different loop-L3 sequences confer different specificities at the “master-nucleotide” position, in each case, through interactions with an unstacked, flipped, and inserted “master nucleotide.” Campagne et al. (2014, 2015) supported this hypothesis by identifying examples of L3 −loop sequences that conferred specificity for cytidine, thymidine, and adenosine at the “master-nucleotide” position, and by providing evidence that swapping L3-loop sequences swaps specificity at the “master-nucleotide” position. Our results provide further support for the hypothesis by identifying an example of an L3-loop sequence, the *Mtb* σ^L^ loop-L3 sequence, that confers specificity for guanosine at the “master-nucleotide” position, and by documenting that specificity for guanosine involves interactions with an unstacked, flipped, and inserted “master nucleotide” (Figs. 3–5, S4A).

In the crystal form identified and analyzed in this work, σR2 of each molecule of transcription initiation complex makes no interactions with other molecules of transcription initiation complex in the crystal lattice (Fig. S7A), and, therefore, with this crystal form, it should be possible to substitute σR2 without losing the ability to form crystals (Fig. S7A). This potentially provides a platform for systematic structural analysis of σR2 and σR2-DNA interactions for the thirteen *Mtb* σ factors, by determination of crystal structures of transcription initiation complexes containing “chimeric σ factors” (see Kumar et al., 1995; Rhodius et al., 2013) comprising σR2 of a *Mtb* σ factor of interest fused to the σR2/4 linker through σR4 of *Mtb* σ^L^ (Fig. S7B; left red arrow) and containing the promoter sequence for the *Mtb* σ factor of interest. In the crystal form identified and analyzed in this work there also are no lattice interactions for the connector between σR2 and σR4, and there likely would be no lattice interactions even if that connector were to contain σR3 and a σR3/4 linker, as in group-1, group-2, and group-3 σ factors (Figure S7A). Accordingly, this crystal form potentially provides a platform for systematic structural analysis not only of σR2 and its protein-DNA interactions, but also of the connector between σR2 and σR4 and its protein-DNA interactions, for the thirteen *Mtb* σ factors, by determination of crystal structures of transcription initiation complexes containing chimeric σ factors comprising σR2 and the connector of one *Mtb* factor fused to σR4 of *Mtb* σ^L^ (Figure S7B, right red arrow).and containing the promoter sequence for the *Mtb* σ factor of interest.

## ACKNOWLEDGEMENTS

This work was supported by NIH grant GM041376 to R.H.E. We thank S. Rodrigue for plasmids and APS at Argonne National Laboratory and Stanford Synchrotron Radiation Lightsource for beamline access.

## AUTHOR CONTRIBUTIONS

W.L. and M.C. prepared RNAP derivatives. W.L., Y.F., and K.D. performed structure determination. W.L., D.D., and S.M. performed sequence analyses and biochemical experiments. R.H.E. designed the study, analyzed data, and wrote the paper.

## STAR METHODS

### *M. tuberculosis* RNAP core enzyme

*Mtb* RNAP core enzyme was prepared by co-expression of genes for *Mtb* RNAP β’ subunit, β subunit, N-terminally decahistidine-tagged α subunit, and ω subunit in *E. coli,* followed by cell lysis, polyethylenimine precipitation, ammonium sulfate precipitation, immobilized-metal-ion affinity chromatography on Ni-NTA agarose (Qiagen), and anion-exchange chromatography on Mono Q (GE Healthcare), as in Lin et al., 2018.

### *M. tuberculosis* RNAP σ^A^

*Mtb* RNAP σ^A^ was prepared by expression of a gene for N-terminally hexahistidine-tagged *Mtb* σ^A^ in *E. coli,* followed by cell lysis, immobilized-metal-ion affinity chromatography on Ni-NTA agarose (Qiagen), and anion-exchange chromatography on Mono Q (GE Healthcare), as in Lin *et al,* 2017.

### *M. tuberculosis* σ^L^

*Mtb* RNAP σ^L^ was prepared by expression of a gene for N-terminally hexahistidine-tagged *Mtb* σ^L^ in *E. coli*, followed by cell lysis, immobilized-metal-ion affinity chromatography on Ni-NTA agarose (Qiagen), and anion-exchange chromatography on Mono Q (GE Healthcare), as follows. *E. coli* strain BL21(DE3) (Invitrogen) was transformed with plasmid pSR32 (Jacques et al., 2006; gift of S. Rodrigue, Universitė de Sherbrooke, Canada), encoding N-terminally hexahistidine-tagged *Mtb* σ^L^ under control of the bacteriophage T7 gene 10 promoter. Single colonies of the resulting transformants were used to inoculate 50 mL LB broth containing 50 μg/ml kanamycin, and cultures were incubated 16 h at 37°C with shaking. Aliquots (10 ml) were used to inoculate 1 L LB broth containing 50 μg/ml kanamycin, cultures were incubated at 37°C with shaking until OD_600_ = 0.8, cultures were induced by addition of isopropyl-β-D-thiogalactoside to 1 mM, and cultures were incubated 16 h at 16°C. Cells were harvested by centrifugation (4,000 x g; 15 min at 4°C), re-suspended in buffer A (10 mM Tris-HCl, pH 7.9, 300 mM NaCl, 5 mM DTT, 0.1 mM phenylmethylsulfonyl fluoride, and 5% glycerol), and lysed using an EmulsiFlex-C5 cell disruptor (Avestin). The lysate was centrifuged (20,000 x g; 30 min at 4°C), the pellet was re-suspended in buffer B (8 M urea, 10 mM Tris-HCl, pH 7.9, 10 mM MgCl_2_, 10 mM ZnCl_2_, 1 mM EDTA, 10 mM DTT and 10% glycerol), and the suspension was further centrifuged (20,000 x g; 30 min at 4°C). The supernatant was loaded onto a 5 ml column of Ni^2+^-NTA-agarose (Qiagen) pre-equilibrated in buffer B, and the column was washed with 9×15 ml buffer B containing 5 mM, 10 mM, 20 mM, 30 mM, 40 mM, 50 mM, 60 mM, 70 mM, and 80 mm imidazole, and eluted with 50 ml buffer B containing 200 mM imidazole. The sample was subjected to step dialysis for renaturation [10 kDa MWCO Amicon Ultra-15 centrifugal ultrafilters (EMD Millipore); dialysis 4 h at 4°C against 8 volumes 50% (v/v) buffer C (10 mM Tris-HCl, pH 7.9, 200 mM NaCl, 1 mM DTT, 0.1 mM EDTA, and 5% glycerol) in buffer B; dialysis 4 h at 4°C against 8 volumes 75% (v/v) buffer C in buffer B; dialysis 4 h at 4°C against 8 volumes 87.5% (v/v) buffer C in buffer B; and dialysis 4 h at 4°C against 8 volumes buffer C]; further purified by gel filtration chromatography on a HiLoad 16/60 Superdex 200 prep grade column (GE Healthcare) in 20 mM Tris-HCl, pH 8.0, 100 mM NaCl, 5 mM MgCl_2_, and 1 mM 2-mercaptoethanol; concentrated to 10 mg/ml in the same buffer using 10 kDa MWCO Amicon Ultra-15 centrifugal ultrafilters (EMD Millipore); and stored in aliquots at −80°C. Yields were ~5mg/l, and purities were ~95%.

Alanine-substituted σ^L^ derivatives were prepared as described for preparation of σ^L^, but using plasmid pSR32 derivatives constructed using site-directed mutagenesis (QuikChange Site-Directed Mutagenesis Kit; Agilent).

Selenomethionine-substituted σ^L^ was prepared as described for preparation of σ^L^, but using culture media and culture procedures as in Stols et al., 2004.

### *M. tuberculosis* RNAP *σ* holoenzyme

*Mtb* RNAP σ^A^ holoenzyme was prepared by co-expression of genes for *Mtb* RNAP β’ subunit, RNAP β subunit, RNAP N-terminally decahistidine-tagged α subunit, and RNAP ω subunit, and N-terminally hexahistidine-tagged σ^A^ in *E. coli*, followed by cell lysis, polyethylenimine precipitation, ammonium sulfate precipitation, immobilized-metal-ion affinity chromatography on Ni-NTA agarose (Qiagen), and anion-exchange chromatography on Mono Q (GE Healthcare), as in Lin et al., 2017.

### *M. tuberculosis* RNAP σ^L^ holoenzyme

*Mtb* RNAP core enzyme and *Mtb* σ^L^ or σ^L^ derivative were incubated in a 1:4 ratio in 20 mM Tris-HCl, pH 8.0, 100 mM NaCl, 5 mM MgCl_2_, and 1 mM 2-mercaptoethanol for 12 h at 4°C. The reaction mixture was applied to a HiLoad 16/60 Superdex S200 column (GE Healthcare) equilibrated in the same buffer, and the column was eluted with 120 ml of the same buffer. Fractions containing *Mtb* RNAP σ^L^ holoenzyme were pooled, concentrated to ~10 mg/ml using 30 kDa MWCO Amicon Ultra-15 centrifugal ultrafilters (EMD Millipore), and stored in aliquots at −80°C.

### Oligonucleotides

Oligodeoxyribonucleotides (Integrated DNA Technologies) and the pentaribonucleotide 5’-CpUpCpGpA-3’ (TriLink) were dissolved in nuclease-free water (Ambion) to 3 mM and were stored at −80°C.

### Nucleic-acid scaffolds

Nucleic-acid scaffolds RPitc5_sp4, RPitc5_sp5, RPitc5_sp6, and RPo_sp6 (sequences in Fig. S2) were prepared as follows: Nontemplate-strand oligodeoxyribonucleotide (0.5 mM), template-strand oligodeoxyribonucleotide (0.55 mM), and, where indicated, pentaribonucleotide (1 mM) in 40μl 20 mM Tris-HCl, pH 8.0, 100 mM NaCl, 5 mM MgCl_2_, and 1 mM 2-mercaptoethanol, were heated 5 min at 95°C, cooled to 25°C in 2°C steps with 1 min per step using a thermal cycler (Applied Biosystems), and stored at −80°C.

### Structure determination: assembly of transcription initiation complexes

Transcription initiation complexes were assembled by mixing 16 μl 50 μM *Mtb* RNAP σ^L^ holoenzyme (in 20 mM Tris-HCl, pH 8.0, 75 mM NaCl, 5 mM MgCl_2_, and 5 mM dithiothreitol) and 4 μl 0.4 mM nucleic-acid scaffold (previous section) in 5 mM Tris-HCl, pH 7.7, 0.2 M NaCl, and 10 mM MgCl2, and incubating 1 h at 25°C.

### Structure determination: crystallization, cryo-cooling and crystal soaking

Robotic crystallization trials were performed for *Mtb* RPitc5-σ^L^_sp6 using a Gryphon liquid handling system (Art Robbins Instruments), commercial screening solutions (Emerald Biosystems, Hampton Research, and Qiagen), and the sitting-drop vapor-diffusion technique (drop: 0.2 μl transcription initiation complex (previous section) plus 0.2 μl screening solution; reservoir: 60 μl screening solution; 22°C). 900 conditions were screened. Under several conditions, *Mtb* RPitc5-σ^L^_sp6 crystals appeared within 2 weeks. Conditions were optimized using the hanging-drop vapor-diffusion technique at 22°C. The optimized conditions for *Mtb* RPitc5-σ^L^_sp6 (drop: 1 μl *Mtb* RPitc5-σ^L^_sp6 in 20 mM Tris-HCl, pH 8.0, 75 mM NaCl, 5 mM MgCl_2_, and 5 mM dithiothreitol plus 1 μl 100 mM sodium citrate, pH 5.5, 200 mM sodium acetate, and 10% PEG4000; reservoir: 400 μl 100 mM sodium citrate, pH 5.5, 200 mM sodium acetate, and 10% PEG4000; 22°C) yielded high-quality, rod-like crystals with dimensions of 0.4 mm x 0.1 mm x 0.1 mm in two weeks (Fig. S2). Crystals were transferred to reservoir solution containing 18% (v/v) (2R,3R)-(-)-2,3-butanediol (Sigma-Aldrich) and flash-cooled with liquid nitrogen. Analogous procedures were used for *Mtb* RPitc5-σ^L^_sp4, RPitc5-σ^L^_sp5, RPitc-σ^L^_sp6, [BrU]RPo-σ^L^_sp6, and [SeMet15,76] RPo-σ^L^_sp6.

### Structure determination: data collection and reduction

Diffraction data were collected from cryo-cooled crystals at Argonne National Laboratory beamline 19ID-D and Stanford Synchrotron Radiation Lightsource SSRL-9-2. Data were processed using HKL2000 (Otwinowski et al., 1997). The resolution cut-off criteria were: (i) I/σ > =1.0, (ii) CC_1/2_ (highest resolution shell) > 0.5.

### Structure determination: structure solution and refinement

The structure of *Mtb* RPitc5-σ^L^_sp6 was solved by molecular replacement with MOLREP (Collaborative Computational Project, 1994) using the structure of *Mtb* RPo (PDB 5UHA; Lin et al., 2017), omitting σ^A^ and nucleic acids, as the search model. One molecule of RNAP was present in the asymmetric unit. Early-stage refinement included rigid-body refinement of RNAP core enzyme, followed by rigid-body refinement of each subunit of RNAP core enzyme, followed by rigid-body refinement of 38 domains of RNAP core enzyme (methods as in Zhang et al., 2012). Electron density for σ^L^ and nucleic acids was unambiguous, but was not included in models in early-stage refinement. Cycles of iterative model building with Coot (Emsley et al., 2010) and refinement with Phenix (Adams et al., 2010)) then were performed. Improvement of the coordinate model resulted in improvement of phasing, and electron density maps for σ^L^ and nucleic acids, which were not included in models at this stage, improved over successive cycles. σ^L^ and nucleic acids then were built into the model and refined in stepwise fashion.

The final model was generated by XYZ-coordinate refinement with secondary-structure restraints, followed by group B-factor and individual B-factor refinement. The final model, refined to R_work_ and R_free_ of 0.19 and 0.23, respectively, was deposited in the PDB with accession code 6DVC (Table 1).

Analogous procedures were used to solve and preliminarily refine structures of *Mtb* RPitc5-σ^L^_sp4, RPitc5-σ^L^_sp5, RPitc5-σ^L^_sp6, and [BrU]RPo-σ^L^_sp6; models of σ^L^ and nucleic acids then were built into mF_o_-DF_c_ difference maps, and additional cycles of refinement and model building were performed. The final models were deposited in the PDB with accession codes 6DV9, 6DVB, and 6DVD (Table 1).

Analogous procedures were used to solve and preliminarily refine the structure of [SeMet15,76]RPo-σ^L^_sp6; selenium anomalous signals then were used to determine positions of σ^L^ SeMet15 and σ^L^ SeMet76, and to confirm the register of σ^L^ protein residues. The final model was deposited in the PDB with accession code 6DVE (Table 1).

### Transcription assays

For transcription experiments in Figs. 6, S1E, and S6, reaction mixtures contained (10 μl): 75 nM *Mtb* RNAP σ^L^ holoenzyme or *Mtb* RNAP σ^L^ holoenzyme derivative, 25 nM DNA fragment P-N25-lac [5’-GCCGCC-3’, followed by positions −100 to −1 of bacteriophage T5 N25 promoter (Kammerer et al., 1986), followed by positions +1 to +9 of *E. coli P-lac* (Dickson et al., 1975), followed by 5’-AGGATCACAATTTCACACAG-3’; prepared by annealing synthetic oligodeoxyribonucleotides, followed by PCR amplification] or DNA fragment P-*sigL-lac* or single-base-pair-substituted derivative thereof [5’-GCCGCC-3’, followed by positions −100 to −1 of *Mtb P-sigL* (Hahn et al., 2005; Dainese et al., 2006; Rodrigue et al., 2007) or single-base-pair-substituted derivative thereof, followed by positions +1 to +9 of *E. coli P-lac* (Dickson et al., 1975), followed by 5’-AGGATCACAATTTCACACAG-3’; prepared by annealing synthetic oligodeoxyribonucleotides, followed by PCR amplification], 100 μM [α^32^P]-UTP (0.03 Bq/fmol), 100 μM ATP, and 100 μM GTP in transcription buffer (40 mM Tris-HCl, pH 8.0, 75 mM NaCl, 5 mM MgCl_2_, 2.5 mM DTT, and 12.5% glycerol). Reaction components other than DNA and nucleotides were pre-incubated 5 min at 22°C; DNA was added and reaction mixtures were incubated 5 min at 37°C; and nucleotides were added and reaction mixtures were further incubated 5 min at 37°C. Reactions were terminated by addition of 2 μl loading buffer (80% formamide, 10 mM EDTA, 0.04% bromophenol blue, and 0.04% xylene cyanol). Products were heated 5 min at 95°C, cooled 5 min on ice, and applied to 16% polyacrylamide (19:1 acrylamide:bisacrylamide, 7M urea) slab gels (Bio-Rad), electrophoresed in TBE (90 mM Tris-borate, pH 8.0, and 0.2 mM EDTA), and analyzed by storage-phosphor scanning (Typhoon: GE Healthcare).

Transcription experiments in Fig. S1F, were performed in the same manner as transcription experiments in Figs. 6, S1E, and S6, but using reaction mixtures containing (10 μl): 600 nM *Mtb* RNAP σ^L^ holoenzyme, 400 nM annealed nontemplate and template strands of nucleic-acid scaffolds RPitc5_sp4, RPitc5_sp5, RPitc_sp6, and RPitc5_sp7 (sequences in Fig. S2 for RPitc5_sp4, RPitc5_sp5, RPitc_sp6; 5’-CGTGTCAGTAAGCTGTCACGGATGCAGG-3’ and 5’-CCTGCATCCGTGAGTCGAGGG-3’ for RPitc5_sp7), 1 mM [α^32^P]-UTP (0.003 Bq/fmol), 1 mM ATP, and 1 mM CTP in transcription buffer.

Transcription experiments in Fig. S1G, were performed in the same manner as transcription experiments in Figs. 6, S1E, and S6, but using reaction mixtures containing (10 μl): 75 nM *Mtb* RNAP σ^L^ holoenzyme, 25 nM annealed nontemplate and template strands of nucleic-acid scaffolds RPitc5_sp4, RPitc5_sp5, RPitc_sp6, and RPitc5_sp7 (sequences as in preceding paragraph), 500 μM GpA (added together with nucleotides), 100 μM [α^32^P]-UTP (0.03 Bq/fmol), and 100 μM CTP in transcription buffer.

Transcription experiments in Fig. S3C (left panel and lanes 1-2 in right panel) were performed in the same manner as transcription experiments in Figs. 6, S1E, and S6, but including 500 μM ApA (TriLink) in reaction mixtures (added together with nucleotides).

### Transcript-release assays

Transcript-release assays (Fig. S3B, lanes 3-4 in right panel) were performed by carrying out transcription experiments with transcription complexes immobilized on streptavidin-coated magnetic beads, dividing reaction mixtures into supernatants and pellets by magnetic partitioning, and analyzing transcripts in supernatants (released transcripts) and pellets (unreleased transcripts) (see Yarnell and Roberts, 1999). Reaction mixtures contained (50 μl): 75 nM *Mtb* RNAP σ^L^ holoenzyme, 25 nM DNA fragment biotin-P-*sigL*-/*lac* immobilized on streptavidin-coated magnetic beads [prepared by mixing 1.25 pmol biotinylated DNA fragment (biotin incorporated at 5’ end of nontemplate-strand oligodeoxyribonucleotide during synthesis) and 0.05 mg Streptavidin MagneSphere Paramagnetic Particles (Promega; pre-washed with 3×150 μl transcription buffer) in 100 μl transcription buffer 30 min at 22°C, and performing three cycles of removal of supernatant by magnetic partitioning followed by re-suspension in 150 μl transcription buffer at 22°C], 500 μM ApA, 100 μM [α^32^P]-UTP (0.03 Bq/fmol), 100 μM ATP, and 100 μM GTP in transcription buffer. Reaction components other than bead-immobilized DNA, ApA, and NTPs were pre-incubated 5 min at 22°C; bead-immobilized DNA was added and reaction mixtures were incubated 5 min at 37°C; and ApA and NTPs were added and incubated 5 min at 37°C. Reaction mixtures were separated into supernatants and pellets by magnetic partitioning. Supernatants were mixed with 10 μl loading buffer, heated 5 min at 95°C, cooled 5 min on ice, and analyzed by urea-PAGE and storage-phosphor imaging as in the preceding section. Pellets were washed with 3×200 μl transcription buffer at 22°C; mixed with 50 μl loading buffer, heated 5 min at 95°C, cooled 5 min on ice, and analyzed by urea-PAGE and storage-phosphor imaging as in the preceding section.

### Data analysis

Data for transcription activities are means of at least two technical replicates.

### Data availability

Atomic coordinates and structure factors for the crystal structures of *Mtb* RPitc5-σ^L^_sp4, RPitc5σ^L^_sp5, RPitc5-σ^L^_sp6, [BrU]RPo-σ^L^_sp6, and [SeMet15,76]RPo-σ^L^_sp6 have been deposited in the PDB with accession codes PDB 6DV9, 6DVB, 6DVC, 6DVD, and 6DVE.

## SUPPLEMENTAL FIGURE LEGENDS

**Figure S1 (related to Fig. 1).**
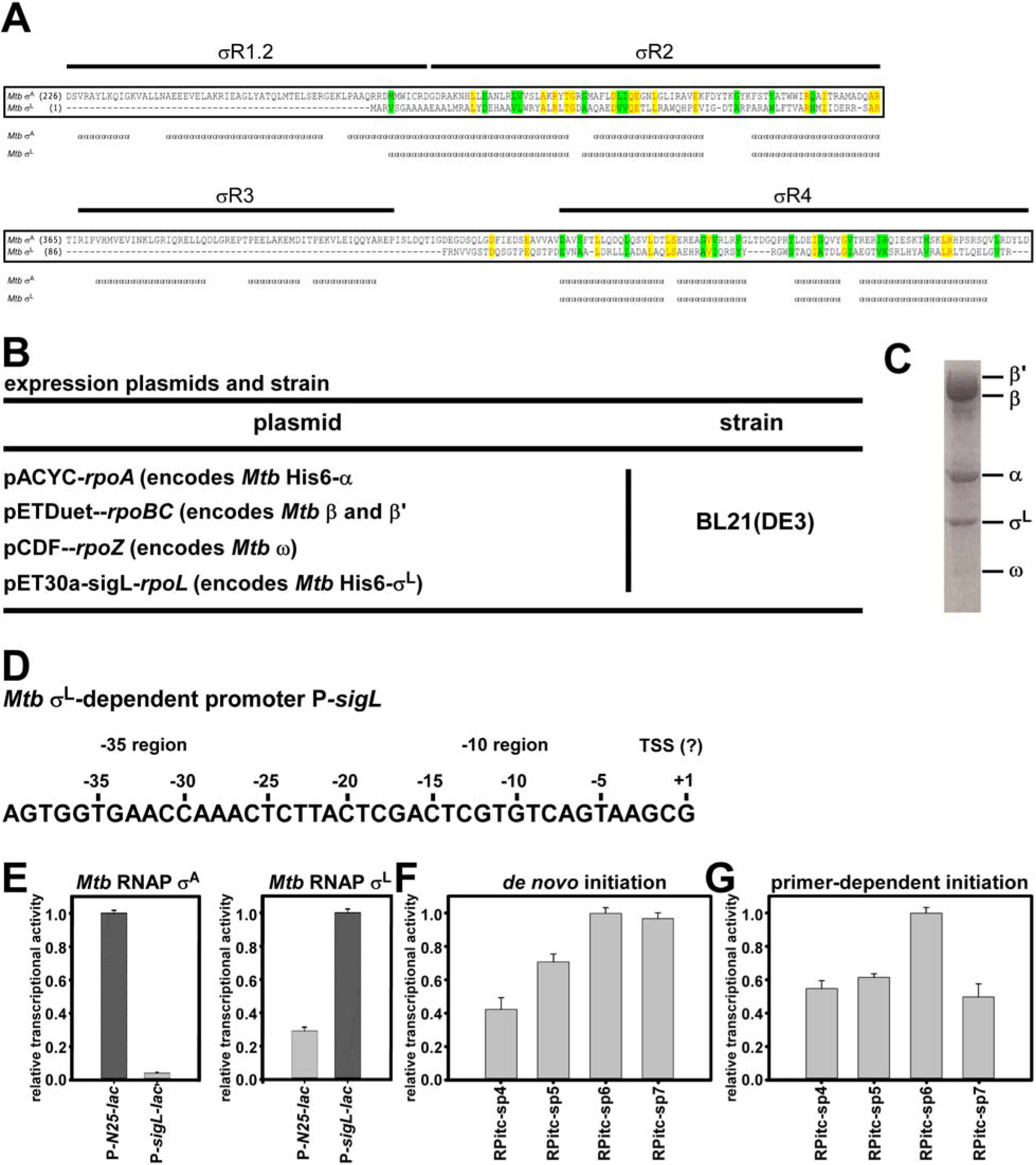
Structure determination: *Mtb* RNAP-σ^L^ holoenzyme and *Mtb* σ^L^-dependent promoter *P-sigL*. **(A)** Sequence alignment of *Mtb* σ^A^ (residues 1 to 225 omitted) and *Mtb* σ^L^. σ conserved regions (σR1.1, σR1.2, σR2, σR3, and σR4) are shown above sequences. Helices, defined from crystal structures in Lin et al., 2017 and in this work (Figs. 1–2), are shown below sequences. **(B)** Plasmids and strain used for production of *Mtb* RNAP-σ^L^ holoenzyme in *E. coli.* **(C)** Coomassie-stained SDS-polyacrylamide gel electrophoresis of *Mtb* RNAP-σ^L^ holoenzyme produced in *E. coli.* **(D)** Sequence of *Mtb* σ^L^-dependent promoter *P-sigL* showing −35 region, −10 region, and reported transcription start site (TSS; Hahn et al., 2005; Dainese et al., 2006; see, however, Rodrigue et al., 2007). **(E)** Transcription experiments demonstrating *Mtb* RNAP σ^A^ holoenzyme selectively recognizes σ^A^-dependent promoter P-N25, and *Mtb* RNAP σ^L^ holoenzyme selectively recognizes σ^L^-dependent promoter P-*sigL.* **(F)** Transcription experiments demonstrating *de novo* transcription initiation by *Mtb* RNAP-σ^L^ holoenzyme on P-sigL derivatives having spacer (“sp”) lengths of 4, 5, 6 and 7 bp (sequences in Fig. S2). **(G)** Transcription experiments demonstrating primer-dependent transcription initiation by *Mtb* RNAP-σ^L^ holoenzyme on P-sigL derivatives having spacer (“sp”) lengths of 4, 5, 6 and 7 bp (sequences in Fig. S2).

**Figure S2 (related to Fig. 1).**
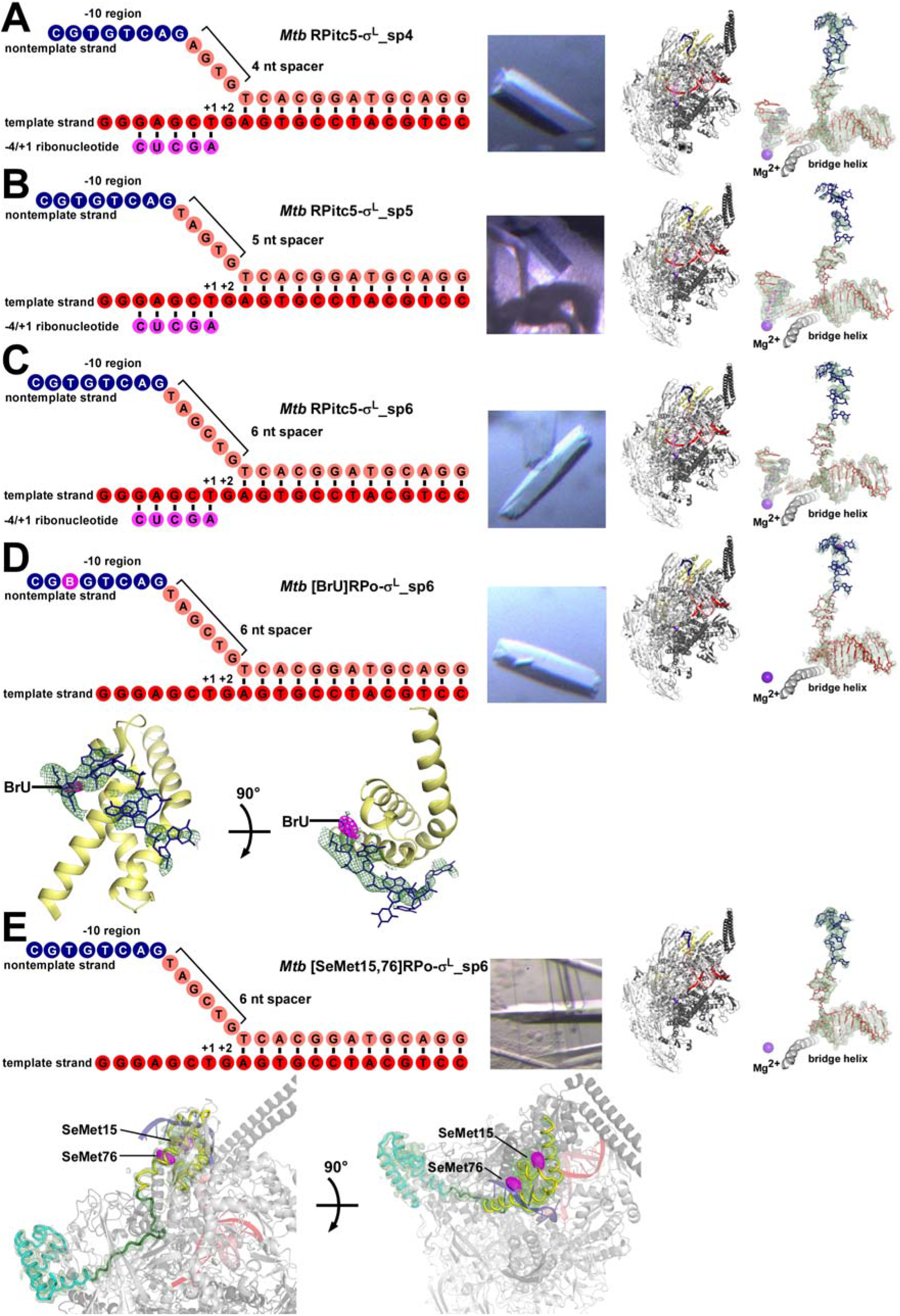
Structure determination: nucleic-acid scaffolds, crystals, and electron densities. **(A)-(C)** Structure determination: *Mtb* −RPitc5-σ^L^_sp4, *Mtb* RPitc5-σ^L^_sp5, and *Mtb* RPitc5-σ^L^_sp6. Left: nucleic-acid scaffold, colored as in Figs. 1–2. Center: crystal. Right: structure and experimental electron density. Green mesh, mF_o_-DF_c_ electron-density omit map (contoured at 2.0σ). **(D)** Structure determination: *Mtb* [BrU]RPo-σ^L^_sp6. Top: as in (A)-(C). Bottom: detail of electron density and Br anomalous difference density for promoter −10 element (two orthogonal views). Green mesh, mF_o_-DF_c_ electron-density omit map (contoured at 2.0σ); magenta mesh, Br anomalous difference density (contoured at 3.0σ). **(E)** Structure determination: *Mtb* [SeMet15,76]RPo-σ^L^_sp6. Top: as in (A)-(D). Bottom: detail of electron density and Se anomalous difference density for σ^L^ (two orthogonal views). Green mesh, mF_o_-DF_c_ electron-density omit map (contoured at 2.0σ); magenta mesh, Se anomalous difference density (contoured at 3.0σ).

**Figure S3 (related to Figs. 3-4).**
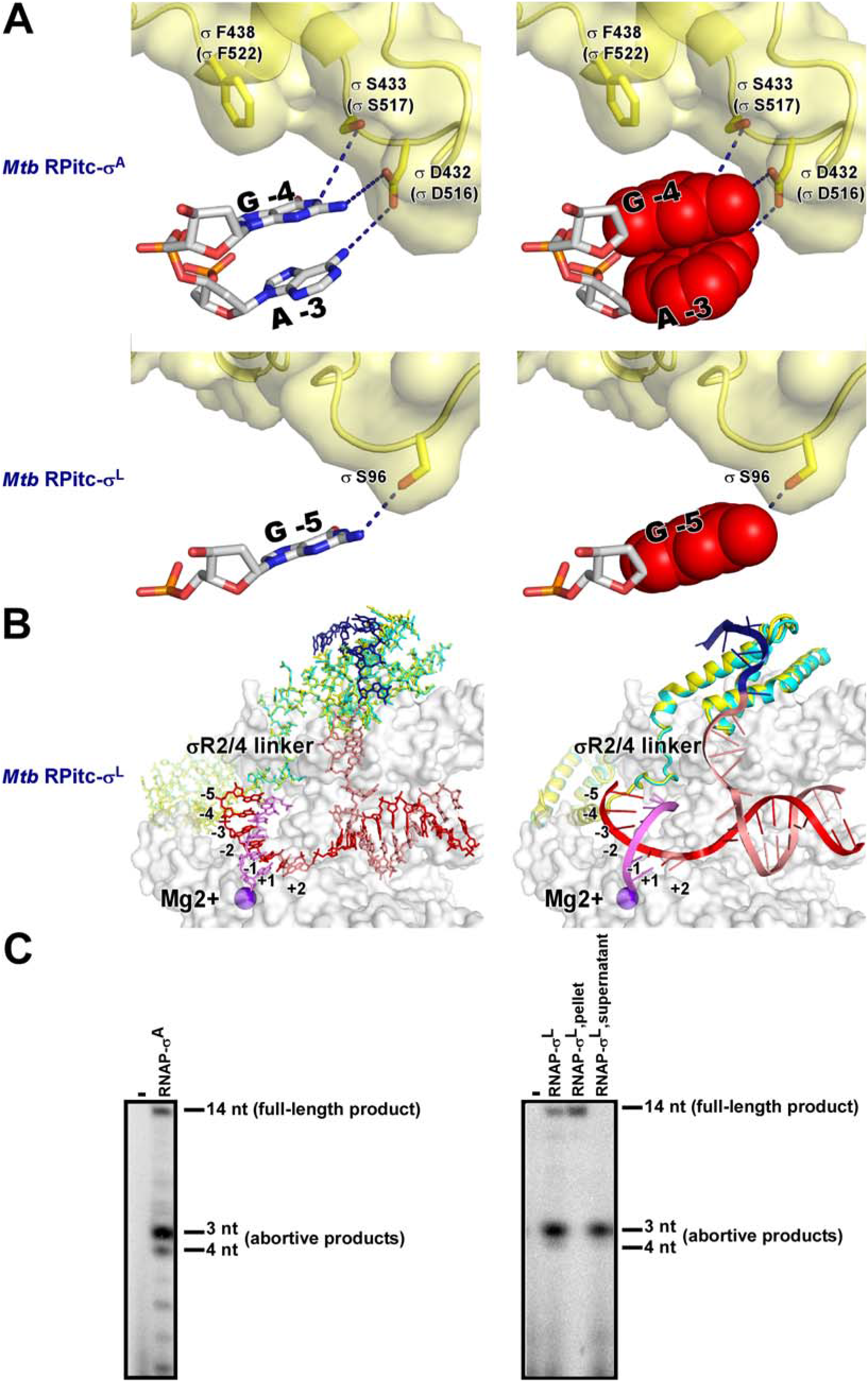
Comparison of protein-nucleic acid interactions with group-1 and ECF σ factors: interactions of σ^A^ σR3/4 linker and σ^L^ σR2/4 linker with transcription-bubble template strand ssDNA. **(A)** Top: interactions of *Mtb* σ^A^ σR3/4 linker with template-strand nucleotides −4 and −3. Bottom: interactions of *Mtb* σ^L^ σR2/4 linker with template-strand nucleotide −5. Red surfaces, space-filling representations of template-strand base moieties. Other colors as in Fig. 5. Residues are numbered as in *Mtb* RNAP, σ^A^, and σ^L^, and, in parentheses, as in *E. coli* RNAP and σ^70^. **(B)** Superimposition of σ^L^ in RPitc (cyan; 5 nt RNA; *Mtb* RPitc5σ^L^_sp6) on σ^L^ in RPo (yellow; 0 nt RNA; *Mtb* [SeMet15,76]RPo-σ^L^_sp6). Left: all-atoms representation of σ^L^; right: ribbon representation of σ^L^. The observation that the conformation of the σ^L^ σR2/4 linker is identical in RPitc5 and RPo indicates that the 5’-end of RNA does not clash with the σ^L^ σR2/4 linker when RNA is ≤5 nt in length. Molecular modeling indicates that the 5’-end of RNA will clash with σ^L^ σR2/4 linker when RNA is >5 nt in length. **(C)** Productive transcription initiation (14 nt RNA products) and abortive transcription initiation (3-4 nt RNA products) by *Mtb* RNAP-σ^A^ holoenzyme and *Mtb* RNAP-σ^L^ holoenzyme. Left panel and lanes 1-2 in right panel show results of transcription experiments; lanes 3-4 in right panel show results of transcript-release experiments (non-released products and released products in lane 3 and 4, respectively). For both *Mtb* RNAP-σ^A^ holoenzyme and *Mtb* RNAP-σ^L^ holoenzyme, the principal abortive products with the analyzed initial-transcribed-sequence are 3 nt and 4 nt in length [ApApU and ApApUpU; identities confirmed by reference to products of parallel reactions omitting ATP and GTP; identities further confirmed by reference to products of parallel reactions with *E. coli* RNAP σ^70^ (see Borowiec and Gralla, 1985)].

**Figure S4 (related to Figs. 3-5).**
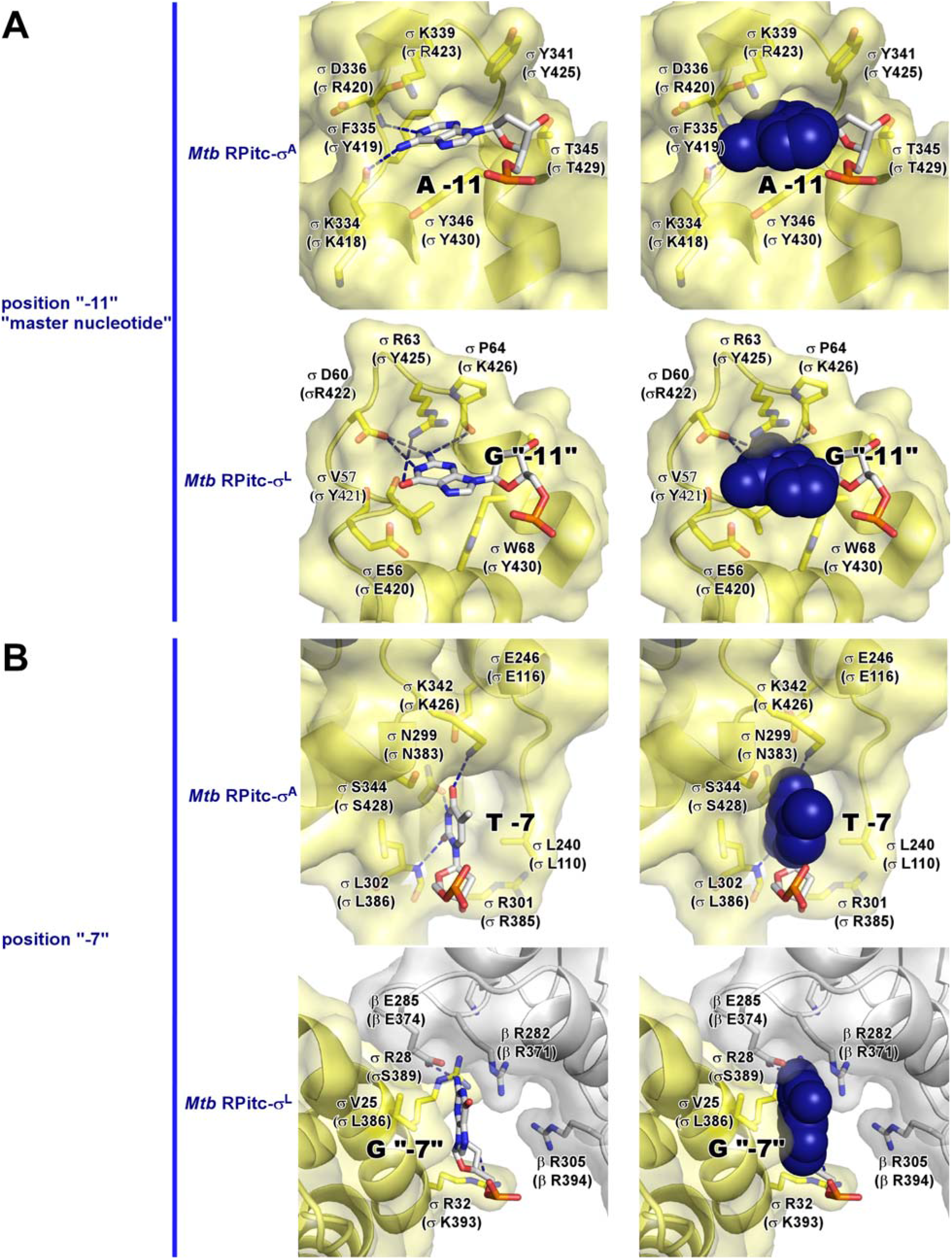
Comparison of protein-nucleic acid interactions with group-1 and ECF σ factors: interactions with unstacked, flipped nucleotides of promoter −10 element inserted into pockets of σR2. **(A)** Interactions of *Mtb* σ^A^ with “master nucleotide” position −11 (top) and *Mtb* σ^L^ with “master nucleotide” position “−11” (bottom). **(B)** Interactions of *Mtb* σ^A^ with position −7 (top) and *Mtb* σ^L^ with position “−7” (bottom). Colors are as in Fig. 5. Residue are numbered as in Fig. S3.

**Figure S5 (related to Figs. 3-4).**
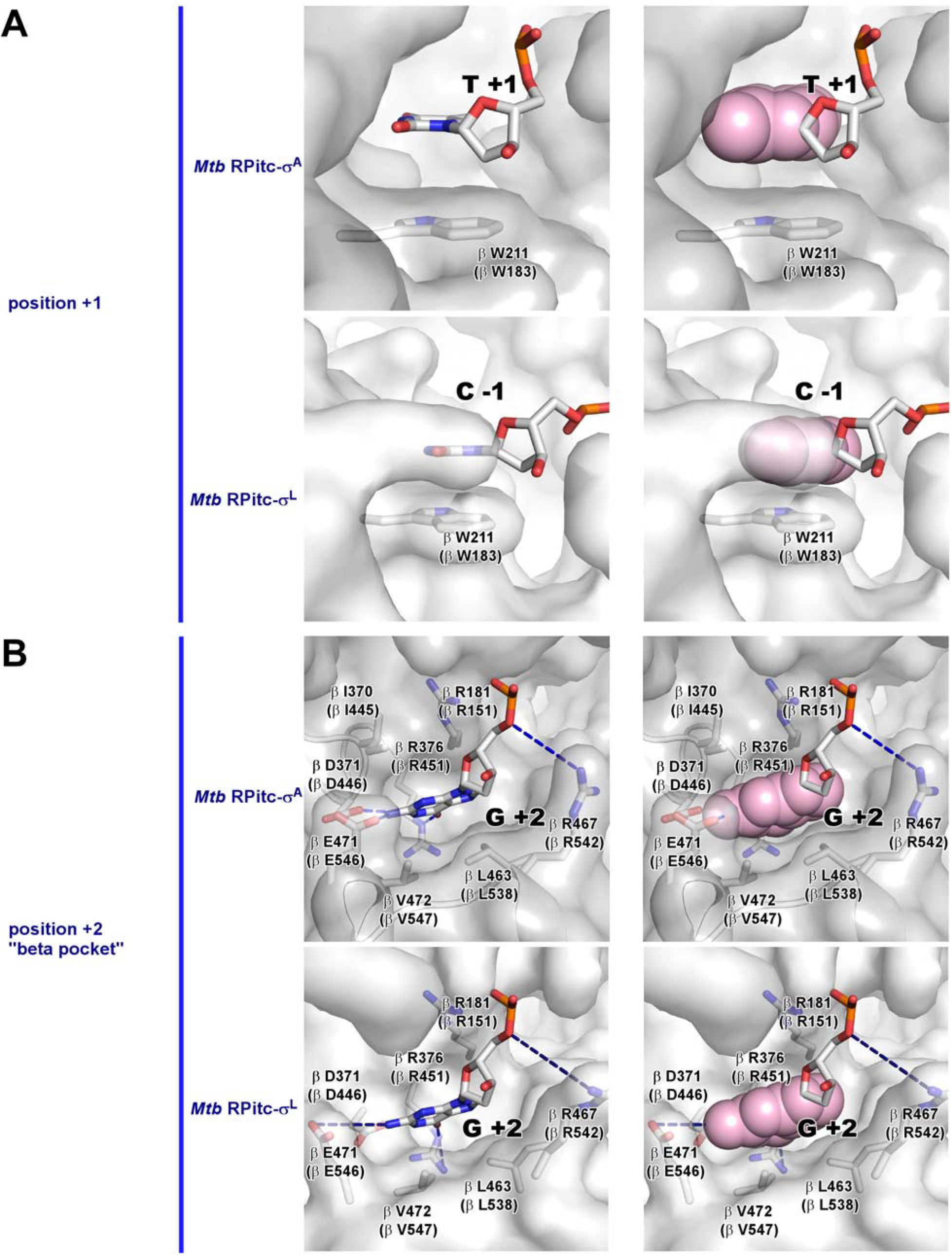
Comparison of protein-nucleic acid interactions with group-1 and ECF σ factors: interactions with promoter core recognition element (CRE). **(A)** Stacking interactions of nontemplate-strand position +1 nucleotide on RNAP β subunit Trp211 in complexes of *Mtb* RPitc-σ^A^ (top) and *Mtb* RPitc-σ^L^ (bottom). **(B)** Unstacking, flipping, and inserting of nontemplate-strand +2 nucleotide into pocket formed by RNAP β subunit (“beta pocket”) in complexes of *Mtb* RPitc-σ^A^ (top) and *Mtb* RPitc-σ^L^ (bottom). Colors are as in Fig. 5. Residue are numbered as in Fig. S3.

**Figure S6 (related to Fig. 6).**
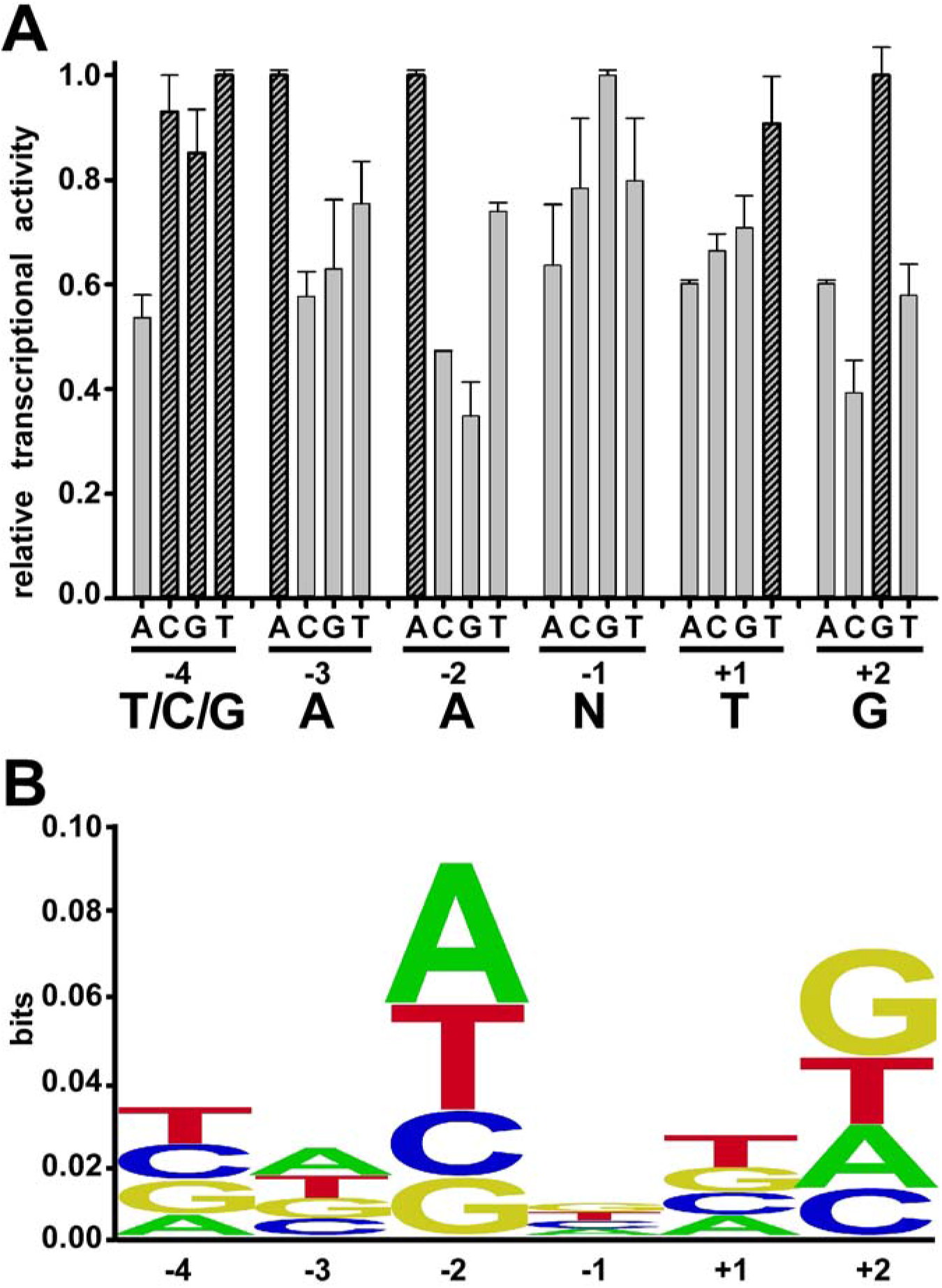
Recognition by *Mtb* σ^L^ of σ^L^-promoter core recognition element (CRE): experimental data. **(A)** Systematic-substitution experiments defining σ^L^-promoter CRE consensus sequence. Relative transcription activities of derivatives of the σ^L^-dependent promoter P-sigL having all possible single-base-pair substitutions at each position of CRE element (positions −4 through +2). Inferred consensus nucleotides are shown at bottom, and data for inferred consensus nucleotides are hatched. **(B)** Sequence logo for σ^L^-promoter CRE consensus sequence [generated using transcription data from (A) and enoLOGOS (Workman et al., 2005; http://biodev.hgen.pitt.edu/enologos/; input setting “energy (2)” and weight type setting “probabilities”).

**Figure S7 (related to Figs. 1-4).**
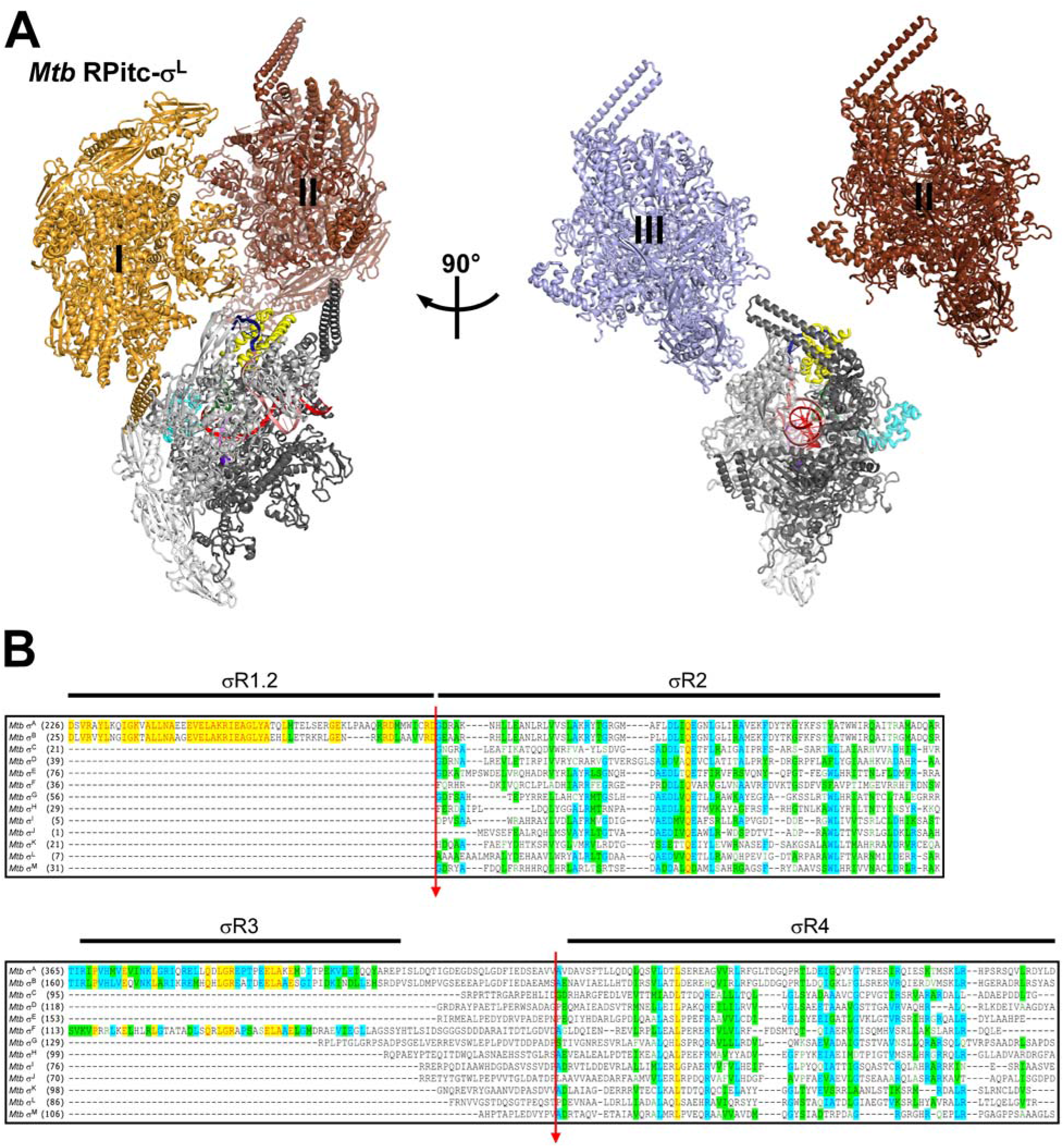
Platform for systematic structural characterization of *Mtb* σ factors. **(A)** Crystal-lattice interactions in crystal form of this work. Figure shows *Mtb* RPitc-σ^L^ (colored as in Figs. 1–4) and the three crystal-lattice neighbors closest to σ^L^ in *Mtb* RPitc-σ^L^ (lattice neighbors I, II, and III in orange, rust, and violet, respectively). Two orthogonal view orientations are shown; for clarity, lattice neighbor III is omitted in first view orientation and lattice neighbor I is omitted in second view orientation. σ^L^ σR2 and σ^L^ σR2/4 linker of *Mtb* RPitc-σ^L^ make no interactions with lattice neighbors. **(B)** Sequence alignment of the 13 *Mtb* σ factors: *Mtb* σ^A^, σ^A^, σ^B^, σ^C^, σ^D^, σ^E^, σ^F^, σ^G^, σ^H^, σ^I^, σ^J^, σ^K^, σ^L^, and σ^M^. Red arrows indicate proposed fusion sites for construction of chimeric σ factors comprising σR1.2-σR2 of *Mtb* σ^A^ through σ^M^ fused to σR2/4 linker-σR4 of *Mtb* σ^L^ (top red arrow) or comprising σR1.2-σR2/4 linker of *Mtb* σ^A^-σ^M^ fused to σR4of *Mtb* σ^L^ (bottom red arrow).

